# SEC-seq reveals translation-focused metabolic strategies for high IgG productivity in clonal CHO cells

**DOI:** 10.64898/2026.04.14.718270

**Authors:** Jasmine Tat, Fides D. Lay, Jennitte Stevens, Nathan E. Lewis

## Abstract

Chinese hamster ovary (CHO) cells are the dominant host for therapeutic protein production, yet intra- and inter-clonal heterogeneity in manufacturing phenotypes, and the underlying metabolic and secretory circuitry, remain poorly defined at single-cell resolution. Here, we apply secretion encoded single-cell sequencing (SEC-seq) to simultaneously measure transcriptomes and secreted IgG in single-cells from a parental production cell line and five CHO clones, each varying in cell-specific productivity. IgG mRNA and recombinant protein secretion are only moderately correlated across single cells, indicating that transcription alone does not explain intra-clonal secretion heterogeneity. By integrating SEC-seq with single-cell metabolic and secretory task scoring, we find that CHO cells accommodating recombinant protein expression burden have more active translation-associated pathways and suppressed energy-intensive endogenous secreted protein processing. Three high-secreting clones converge on this translation-focused state but differ in their subpopulation composition and energy/redox programs coupled to IgG output: one highly productive clone shows a low-growth, glycolytic, NAD/one-carbon–associated and UPR-activated program; a second shows increased oxidative phosphorylation and fatty-acid β-oxidation, and a third shows higher lipid-uptake with modest central carbon metabolism. Genes such as *Aldoa, Ndufab1, Acsl5, Mthfd2* showed clone-specific correlations with IgG, linking glycolysis, mitochondrial respiration, fatty-acid metabolism, and redox to secretion. Together, these results demonstrate that SEC-seq can resolve IgG-coupled metabolic-secretory wiring within and between CHO clones, providing a framework to identify subpopulation and circuit features to engineer or select for improved recombinant protein production.

## Introduction

The biopharmaceutical industry heavily relies on Chinese hamster ovary (CHO) cells, which is the host expression system for 84% of the products in the $115 billion therapeutic monoclonal antibody industry^1^. CHO cells have become the gold standard mammalian host in part due to their inherent genomic plasticity, which has enabled the derivation of highly-productive phenotypes suitable for large-scale manufacturing^2–4^. However, this genomic plasticity in CHO underlies inherent heterogeneity and instability in CHO that is not well characterized and varies across cell lines and process conditions ^4–6^. Thus, despite their widespread use, CHO cells remain “black-box” secretory factories. As the industry is facing the challenge of producing increasingly diverse and complex biologics, we need to identify cellular processes driving recombinant protein production to enable more precise engineering of CHO cells^7–10^.

High-resolution omics methods can provide insights into the cellular programs that drive secretory heterogeneity across cell populations, especially using next-generation sequencing, microfluidics, and single-cell omics. Single-cell transcriptomics studies in CHO have reported relatively low intra-cell line transcriptomic heterogeneity but have revealed subpopulation markers and trajectories associated with instability and productivity^11–14^. Beyond transcriptomics, single-cell imaging-based approaches, including fluorescent microscopy and pulse-chase assays, have been used to probe intracellular IgG dynamics and identify potential secretory pathway bottlenecks^15,16^, and nanovial-based cytometric platforms have enabled sorting and recovery of high-secreting subpopulations. Additionally, nanofluidic and fluorescence-based methods have allowed manipulation, culture, and phenotypic characterization of individual CHO cells with respect to growth, morphology, and secretion dynamics^17,18^. However, most existing single-cell studies in CHO either link host gene expression to bulk phenotypes or recombinant transcription, or interrogate secretion-related phenotypes without direct access to the underlying molecular regulatory state. As a result, the molecular programs that directly govern single-cell secretory capacity remain poorly resolved. To address this gap, we apply SEC-seq^19,20^, a multi-omics approach in which single cells are loaded onto functionalized nanovials to capture secreted IgG, quantify secretion, and concurrently profile the transcriptome, thereby linking gene expression to IgG secretion in CHO cells at single-cell resolution.

Here we apply SEC-seq to probe intra- and inter-clonal transcriptomic heterogeneity coupled to IgG secretion in manufacturing CHO cell lines. We profiled a parental pool and 5 derivative IgG-producing clones spanning a 4-fold range in cell-specific productivity during a fed-batch culture, where IgG transcripts constituted a substantial fraction (up to 40%) of total mRNA transcripts in each cell and impose high transcription, translation, and secretory burden. SEC-seq reveals that clonal CHO cells respond to this burden by adopting a conserved translation-focused state, in which translation-centric pathways are upregulated and high-cost secretory and protein-processing functions are selectively suppressed. High-secreting clones further converge on this translation-optimized program but differ in how glycolysis, oxidative phosphorylation, fatty-acid metabolism, and UPR signaling are wired to sustain high IgG output, as revealed by pathway- and gene-level correlations with IgG across distinct transcriptomic subpopulations. Thus, there are transcriptomic subpopulation signatures associated with both shared and clone-specific strategies in CHO manufacturing lines under bioprocess-relevant conditions, providing insights for precision engineering of CHO cells for biologics production.

## Results

### SEC-seq quantifies single-cell transcriptome and IgG secretion in manufacturing CHO cells

SEC-seq uses gelatin coated nanovials to load individual cells and capture single-cell protein secretions using embedded antibodies. These nanovials are compatible with flow cytometry sorters and 10X Chromium-based NGS library preparation^19,20^ (**Figure 1a**). We evaluated SEC-seq compatibility with industrial suspension-based CHO cell lines, for which nanovial loading efficiency may be limited by cell adhesion to gelatin, a process dependent on cell surface receptor (e.g., integrin) expression. We first confirmed that suspension CHO cells could be single-cell loaded into gelatin nanovials and identified using a viability dye (**Figure 1b**). Next, we observed that nanovials modified with an antibody configuration can capture a dynamic and representative range of IgG concentrations (**Figure 1c,d**). Finally, we deployed the complete SEC-seq workflow using a panel of 5 single-cell-derived CHO cell lines (hereon referred to as clones) cultivated in a small-scale fed-batch process. The clones spanned 4-fold in IgG cell-specific productivity in bulk. We also assayed the parental pool cell line (hereon referred to as pool), along with a null-transfected wild-type CHO host as a non-secreting control cell line (**Figure 1e-h**). We confirmed that oligo-barcoded-streptavidin enables the computational deconvolution of nanovial-loaded single-cells and free cells (**Figure 1f**). We quantified the secreted IgG using the SEC-seq feature barcodes, and found the barcodes correlated highly with the FACS-based fluorescent signal and the cell-specific productivity observed in the bulk population (**Figure 1g,h**). This demonstrated that SEC-seq can reproduce the relative ranking of the clone productivity profiles without negatively impacting the sequencing depth and quality of cells in nanovials (**Supplementary Figure 1b-e**). Overall, we found that SEC-seq is compatible with suspension-based CHO cell lines, allowing us to link the single-cell transcriptome with IgG secretion.

**Figure 1.**
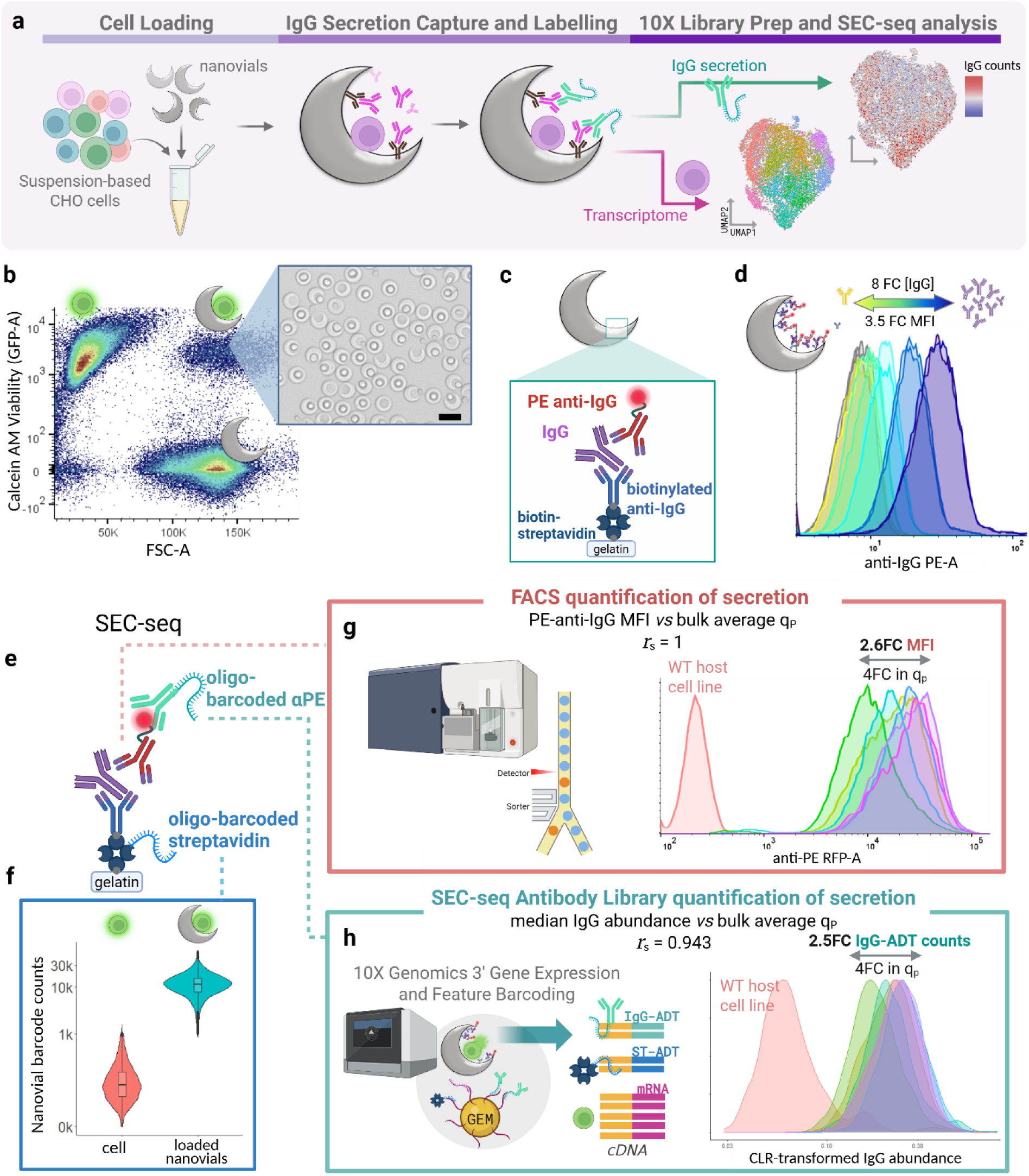
SEC-seq workflow for simultaneous quantification of single-cell transcriptomes and IgG secretion in CHO cells. (**a**) Schematic overview of the SEC-seq workflow, enabling quantification of single-cell transcriptome and IgG secretion. Single cells are first loaded into biotinylated gelatin-coated nanovials that are subsequently functionalized with anti-IgG antibodies to capture secreted IgG. Oligo-barcoded antibodies enable NGS quantification of captured IgG. Nanovials and oligo-barcoded antibodies are compatible with 10x Genomics 3’ Gene Expression and Feature Barcoding protocols, allowing us to generate single-cell transcriptome and IgG secretion libraries. (**b**) Calcein AM viability staining combined with forward scatter (FSC) enables identification and sorting of cell-loaded nanovials. Brightfield image shows GFP+ and FSC+ sorted cell-loaded nanovials. Scale bar: 50 µm (**c**) Schematic of nanovial modification for IgG capture and detection. Biotinylated gelatin nanovials are first modified with streptavidin. Biotinylated anti-IgG antibodies are then added enabling secreted IgG capture. Phycoerythrin (PE) conjugated anti-IgG antibodies are added to detect captured IgG within nanovial cavities. (**d**) Detection of IgG across an 8-fold concentration range using the configuration in (**c**) and incubation with serially diluted crude harvest supernatant. (**e**) Antibody configuration used for SEC-seq. (**f**) Oligo-barcoded streptavidin enables classification of free-cells and cell-loaded nanovials through the feature barcode library. (**g**) PE-conjugated anti-IgG antibody enables fluorescent-based quantification of secreted IgG in cell-loaded nanovials. Density plots show PE fluorescence intensity across non-secreting host cell line and clonal cell lines spanning 4 fold-change in cell specific productivity (q_p_). Spearman correlation coefficient (*r_s_*) is calculated for the PE-A median fluorescence intensity (MFI) and bulk population average q_p_. (**h**) SEC-seq samples from (**g**) are then processed using 10X Genomics 3’ Gene Expression and Feature Barcoding protocols to generate cDNA libraries for single-cell transcriptome, oligo-barcoded streptavidin, and oligo-barcoded anti-PE antibodies. Density plots show centered-log-ratio-(CLR) normalized IgG abundance for each cell line sample. Spearman correlation coefficient (*r_s_*) is similarly calculated for the SEC-seq median abundance and bulk population average q_p_. Schematics in **a, c, d, e, f, h** created with BioRender.com

### SEC-seq reveals clone-level coupling between IgG transcription and secretion

To investigate how transcriptomic features couple with IgG secretion in clonally derived CHO cell lines, we applied SEC-seq to a panel of production clones exhibiting up to a 4-fold difference in cell-specific productivity (qp) along with their parental pool, on Day 8 of a small-scale fed-batch bioreactor culture, representing a highly productive metabolic phase (**Figure 2a,b**). SEC-seq resolved significant differences in secreted IgG counts between low and high qp clones (**Figure ^2^c**) and reproduced the expected bulk IgG productivity phenotypes (**Figure 1h**), confirming its ability to capture secretion heterogeneity.

**Figure 2.**
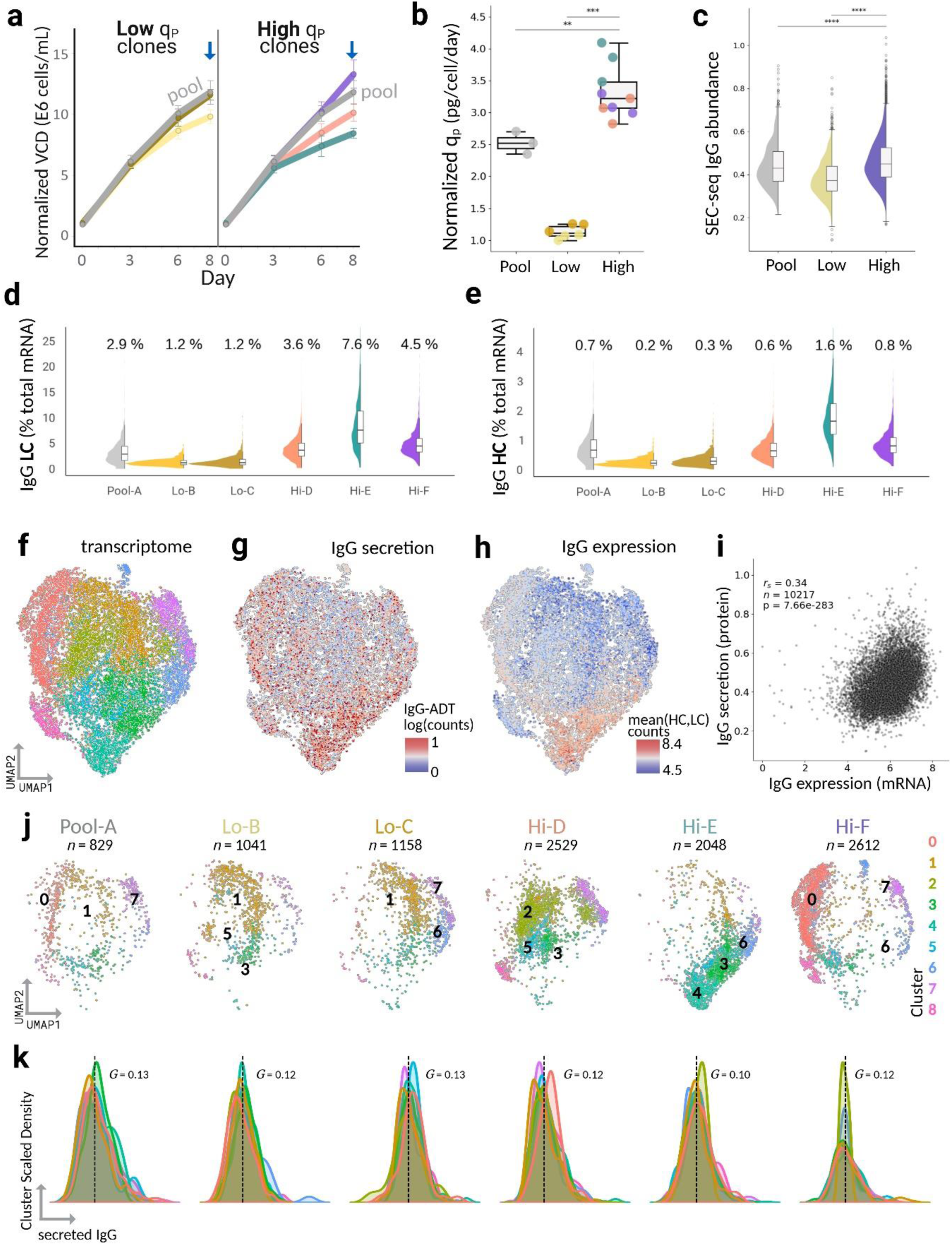
SEC-seq quantifying transcriptomic and IgG secretion profiles for low and high-secreting CHO clones. (**a**) Normalized viable cell density (VCD) growth profiles over a fed-batch process for low- and high-cell specific productivity (q_p_) clones, over a small-scale fed-batch process, with the parental pool included in both panels (grey). Day 8 SEC-seq sampling is indicated (blue arrow). VCD was normalized against Day 0 inoculation cell concentration. (**b**) Normalized q_p_ (pg/cell/day) on Day 8 for the parental pool, Low q_p_ clones, and High q_p_ clones. Colors denote individual clones. q_p_ was normalized against the lowest value. Mann-Whitney U one-sided tests confirm greater median q_p_ for the parental pool and High q_p_ clones relative to the Low q_p_ clones (\*\**p*<0.01; \*\*\**p*<0.001). (**c**) SEC-seq centered log-ratio (CLR)-transformed IgG abundance (oligo barcode counts) for the parental pool, Low q_p_ clones and High q_p_ clones. Mann-Whitney U one-sided tests confirm greater SEC-seq counts for the Pool and High clones relative to the Low clones (\*\**p*<0.01; \*\*\**p*<0.001). (**d,e**) Proportion of transcripts assigned to IgG light chain (LC) and heavy chain (HC) respectively for each cell line sample. (**f**) UMAP projection of single-cell transcriptomes data colored by Louvain clusters (n=9). (**g**) UMAP overlaid with SEC-seq single-cell IgG secretion. (**h**) UMAP overlaid with IgG mRNA expression (mean expression of LC and HC) (**i**) Scatterplot of single-cell IgG expression (mRNA) versus SEC-seq quantified IgG secretion (protein abundance) across all cells, with Spearman rho (*r_s_*), number of cells (*n*), and p-value annotated. (j) UMAP split by cell line sample in each panel colored by Louvain clusters with the top 3 abundant clusters labelled. “Lo-” indicates a low q_p_ clone and “Hi-“ indicates a high q_p_ clone. Number indicates the number of single cells (loaded in nanovial) per cell line sample. (**k**) Density plots of IgG secretion per cluster per cell line sample, with median cell line sample IgG (dotted line) and median bootstrapped Gini coefficient (*G*) annotated. All significance values reflect Benjamini–Hochberg–adjusted p-values.

Consistent with previous bulk transcriptome analyses of CHO cell lines expressing heterologous proteins^9,21,22^, we observed that IgG expression imposes a substantial transcriptional burden on CHO clones. In the highest-producing clone (Hi-E), IgG transcripts accounted for a median of 1.6% and 7.6% of the transcriptome, with maximal values reaching 6.9% and 39.6% for IgG heavy chain (HC) and light chain (LC) respectively (**Figure 2d,e**, **Supplementary Figure 1j**). In contrast, highly-expressed housekeeping genes accounted for a small fraction of the transcriptome. Across the top-expressed housekeeping genes, the median per-gene transcriptome proportion was <0.1%, while maximal per-gene proportions reached up to 1.4% (**Supplementary Figure 3h,i**).

SEC-seq enabled simultaneous profiling of both transcriptomes and secreted IgG for individual cells across all cultures (**Figure 2f–h**). We first examined the relationship between IgG transcription and secretion across the entire population. Single-cell IgG expression, quantified as the average expression of heavy- and light-chain genes, was moderately correlated with IgG secretion (Spearman *rs* = 0.34, *p* = 7.66 × 10⁻²⁸³; **Figure 2i**). However, for individual cell lines, this correlation was weak (**Supplementary Figure 1i**), indicating that IgG transcription and secretion are coupled at the population level but weakly at the single-cell level.

UMAP projections and clustering of the transcriptome revealed that each cell line is predominantly composed of 1 to 3 clusters (**Figure 2j, Supplementary Figure 2c**). While the low-secreting clones exhibited similar transcriptomic organization, each high-secreting clone contained distinct cluster compositions, highlighting clonal variation. Despite this transcriptomic heterogeneity, the Gini coefficients of secreted IgG across both the whole population and within clusters were uniformly low across all cell lines (**Figure 2k**, **Supplementary Figure 2d–f**). This suggests that secreted IgG levels are evenly distributed among clusters, and secretion is not enriched in specific transcriptomic subpopulations. Similarly, while LC and HC transcript levels varied substantially between clones (**Supplementary Figure 2l,m**), their intra-clone expression distributions were highly uniform (**Supplementary Figure 2e,f**). LC and HC transcript levels exhibited clearer cluster-associated patterns (**Supplementary Figure 2b**), but these differences did not translate into heterogeneity in IgG secretion (**Supplementary Figure 2b**).

### Translation machinery increases with IgG production while high-cost secreted proteins are suppressed

We examined how CHO cells accommodate the high transcriptional and secretory load imposed by recombinant IgG expression. To anchor these analyses, we used a panel of housekeeping genes, expected to maintain stable cellular functions in CHO cells^23^ (**Figure ^3^a**) while analyzing the impact of IgG secretion on core cellular machinery. Across single cells, expression of low-variability housekeeping genes (“Gini genes”) showed minimal association with IgG levels, as reflected by median Spearman correlations close to zero (−0.04 to −0.03 for IgG LC and HC transcript levels and -0.01 for IgG protein levels; **Figure 3b, Supplementary Table 1**). Consistently, single-cell expression distributions were similar between the highest (90th percentile) and lowest (10th percentile) IgG-secreting subpopulations within low and high secreting clones and the parental pool (ANOVA p = 0.216; **Figure 3g, Supplementary Figure 3a-c**), indicating that core cellular functions remain stable despite large shifts in IgG transcript and protein levels.

**Figure 3.**
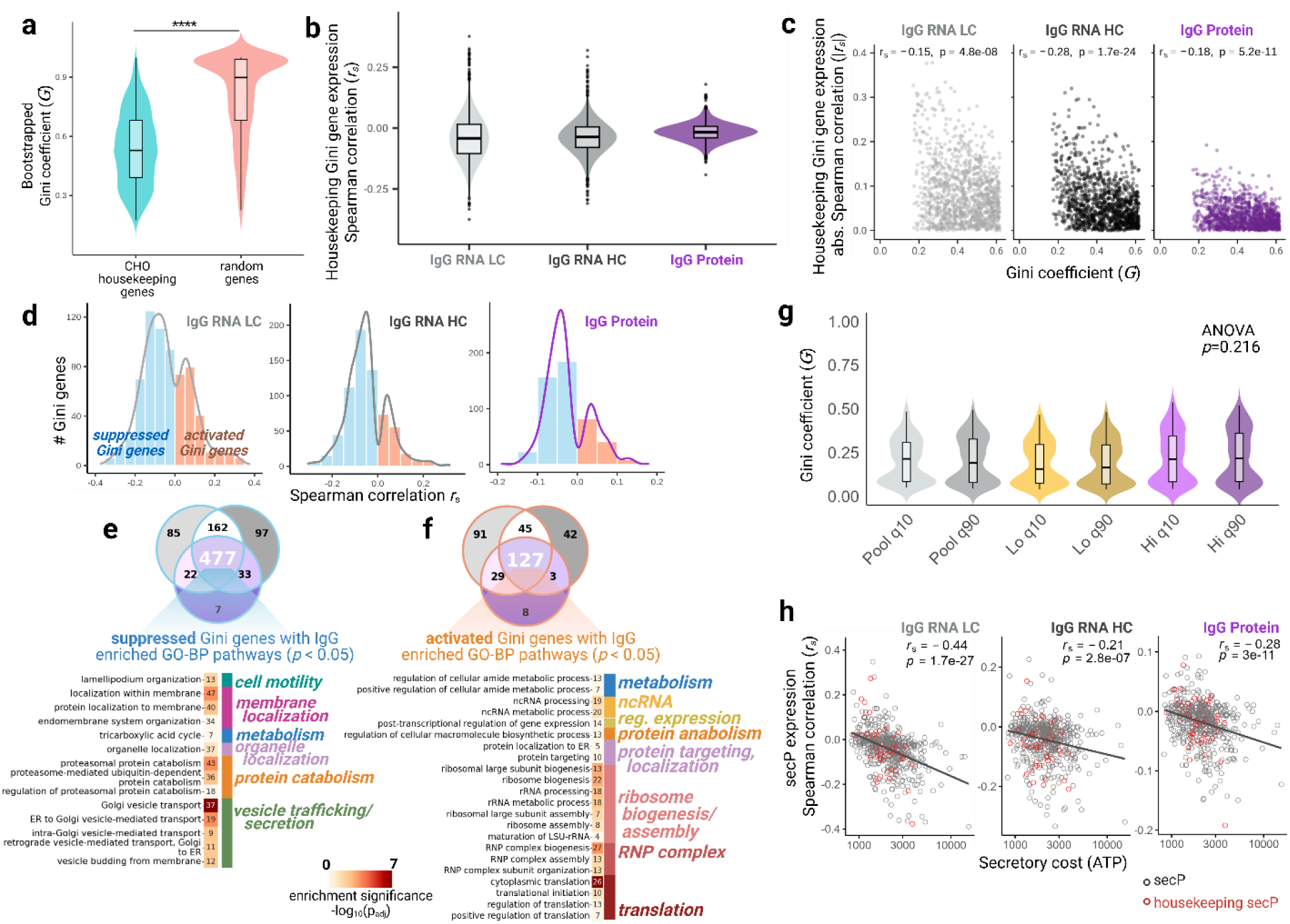
Increasing single-cell IgG RNA and protein levels is associated with activation of translation machinery and suppression of high-cost secretory protein expression. (a) CHO housekeeping genes^23^ (N=1,891) exhibit significantly lower Gini coefficients (*G*) than random genes. Differences were evaluated by bootstrap confidence intervals (5,000 replicates) and permutation tests (10^3^ label shuffles) (\*\*\*\**p*<0.0001). **(b)** Low variability housekeeping genes, or Gini genes (N=1,254), are defined as housekeeping genes in the lowest 20th percentile of overall transcriptome Gini coefficients (random geneset group) from the bootstrapping analysis. Distributions of Spearman correlations (*r_s_*) between Gini gene expression and IgG LC, HC, and secreted protein (partial correlations controlling for sequencing depth for LC and HC analysis). **(c)** Relationship between Gini gene Gini coefficients and absolute correlation with IgG levels. **(d)** Distributions of significant correlations (adjusted *p* < 0.05) for Gini genes with IgG RNA and protein levels. Suppressed (*r_s_* < 0) and activated (*r_s_* > 0) Gini genes are specified as blue and orange, respectively. **(e)** Venn diagram showing overlap of Gini genes that are negatively associated with IgG LC, HC, and protein, as indicated by light grey, dark grey, and purple sets, respectively. GO-BP terms enriched (adjusted *p* < 0.05) among the 477 shared suppressed genes are visualized in a heatmap sorted by pathway theme, with annotations indicating the number of differentially expressed genes mapped to each pathway. Number of Gini genes are annotated per GO-BP term in each heatmap cell. **(f)** Venn diagram of Gini genes positively associated with IgG RNA and protein. Enriched GO-BP terms for 127 shared activated genes are similarly visualized. **(g)** Gini coefficients of Gini housekeeping genes remain statistically similar (ANOVA *p*>0.05) between intra-cell line group low (q10 = 10^th^ percentile of IgG secretion) and high (q90 = 90^th^ percentile of IgG secretion) IgG-secreting subpopulations. (**h**) Increasing IgG RNA and protein levels per cell is associated with lower expression of secreted proteins (secP) with higher ATP cost. Scatterplot of CHO secP (N=563 genes) secretory ATP cost plotted against Spearman correlation coefficient (*r_s_*) of secP expression (CPM) to IgG RNA LC, HC, and Protein levels across all cell line samples. secP genes also classified as housekeeping genes are highlighted in red. All significance values reflect Benjamini–Hochberg–adjusted p-values.

Despite this global stability, subsets of Gini genes exhibited coordinated expression shifts with IgG RNA and protein levels, with a majority of genes identified as significantly anti-correlated (**Figure 3b-d**). Specifically, 477 Gini genes were significantly anti-correlated (“suppressed”; Spearman correlation *rs* < 0, adjusted *p* < 0.05) and 127 positively correlated (“activated” ; Spearman correlation *rs* > 0, adjusted *p* < 0.05) with single-cell IgG transcript and protein abundance (**Figure 3d-f, Supplementary Table 1**). Pathway enrichment revealed that Gini genes involved in translation, ribosome biogenesis and assembly, amide metabolism, protein targeting, and protein anabolism were activated with increased IgG abundance, whereas Gini genes involved in cell motility, membrane and organelle localization, vesicle trafficking and secretion, TCA cycle, and protein catabolism were suppressed (**Figure 3e,f, Supplementary Table 2**). These patterns suggest proportional activated translational capacity alongside suppression of energy-intensive secretory functions to support IgG production. Suppressed vesicle trafficking and secretory functions may be attributed to reduced processing of endogenous secretory proteins that are not critical for cell survival^21,22,24^. To test this, expression of endogenous CHO secreted proteins (secPs) was examined as a function of their ATP biosynthetic costs^24^ (**Supplementary Figure 3f,g, Supplementary Table 3**). Expression of secPs, a subset of which have housekeeping function, decreased with increasing single-cell IgG RNA and protein levels (**Figure 3h**), supporting that CHO cells downregulate non-essential protein synthesis and secretion to conserve energetic resources for recombinant IgG production. Together, these findings suggest that while overall housekeeping stability is maintained, CHO cells fine-tune subsets of core genes to activate translation-associated pathways and suppress the processing of energetically-expensive secretory proteins in response to IgG production burden.

### High-secreting clones coordinate metabolic, secretory, and translation machinery

Given that all clones exhibit a translation-activated and secretion-suppressed response to recombinant IgG burden within their single-cell population, we next investigated how high-secreting clones (Hi-D/E/F) achieve further gains in productivity. Characterization of bulk phenotypes during fed-batch demonstrated that these clones differ not only from low-secreting clones in productivity but also in growth and metabolic behavior: the three high-secreting clones exhibited distinct growth profiles (**Figure 4a**) and overall metabolite consumption rates (**Figure 4b, Supplementary Table 4**). Together with the distinct transcriptomic clustering observed for each high-secreting clone (**Figure 2j**), this suggests that high producers adopt different metabolic–secretory strategies. We therefore leveraged SEC-seq to dissect both inter- and intra-clonal metabolic phenotypes and interrogate how these are coupled to IgG secretion.

**Figure 4.**
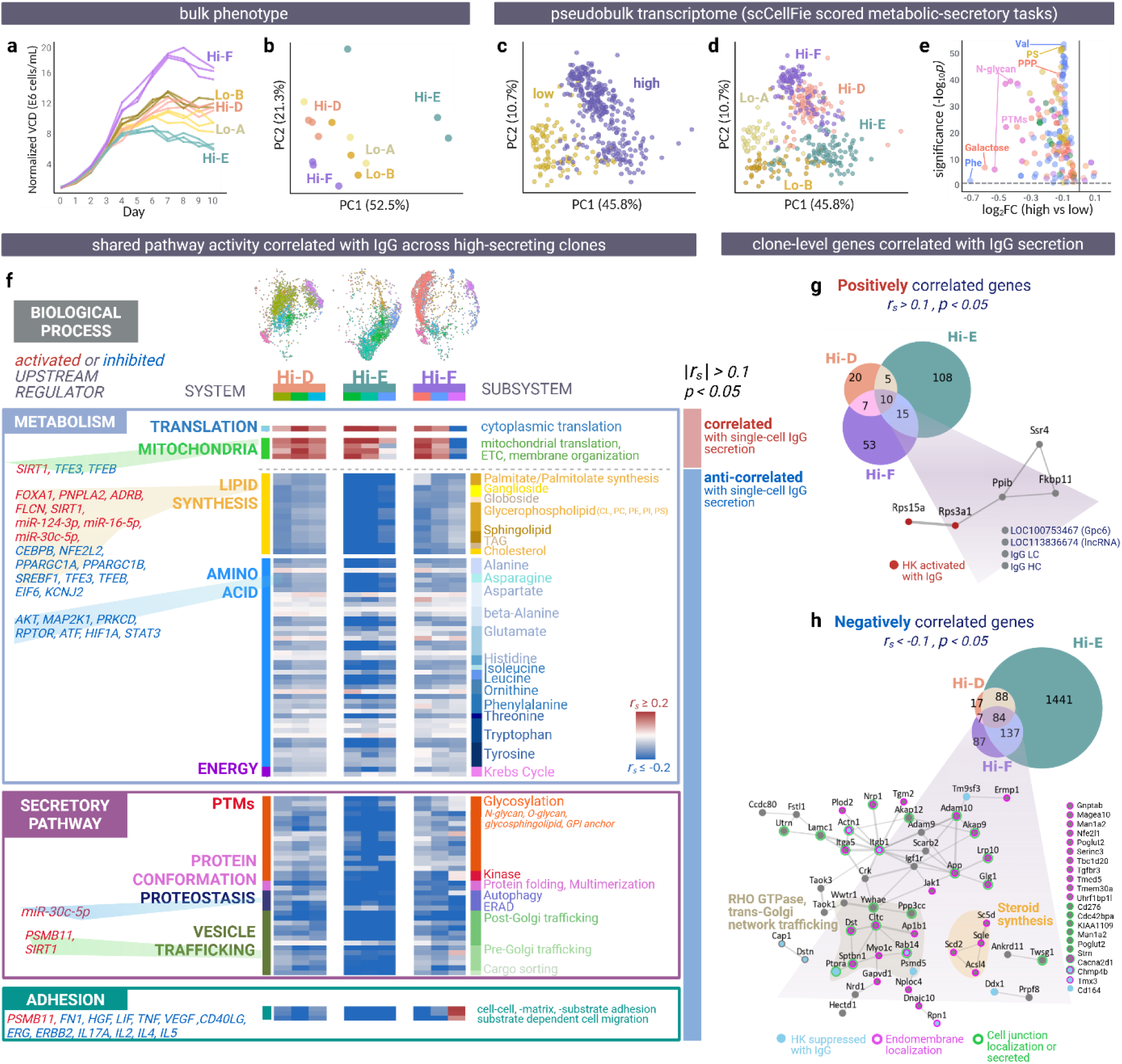
High-secreting clones share common profiles in amino acid metabolism, lipid metabolism, and early to late secretory machinery coupled with single-cell IgG secretion. (**a**) Growth curves (triplicate cultures) for all cell lines over a 10-day fed-batch process; Viable cell density (VCD) normalized to inoculation Day 0 VCD (**b**) Principal component analysis (PCA) of metabolite and amino-acid consumption rates during fed-batch production phase (Days 6–10), colored by cell line. PC1 and PC2 represent the major axes of variation, with the respective percentage variation explained labeled. **(c,d**) PCA of scCellFie metabolic-secretory task scores for pseudobulked average cell groups for each clone-cluster overlaid with (**c**) low- and high-secreting phenotype and (**d**) clone. (**e**) Volcano plot for significant differential task activities (adjusted *p*<0.05) for high- vs low-secreting clone pseudobulk groups overlaid with metabolic-secretory system. The top 4 differentially represented subsystems (for decreasing log_2_FC and decreasing significance) are annotated. (Phe: Phenylalanine metabolism, Val: Valine metabolism, PTMs: post-translational modifications, PS: phosphatidylserine metabolism, PPP: pentose phosphate pathway) (**f**) Heatmap of pathway activities significantly coupled with IgG secretion (|*r_s_* |>0.1, adjusted *p*<0.05, relative to the low-secreting clones) in at least one major cluster from all 3 high-secreting clones. The top 3 abundant clusters per high-secreting clone are considered. Shared upstream regulators activated (red) or inhibited (blue) in at least one clone are annotated. Pathways are organized by major biological process themes: Metabolism, Secretory Pathway, and Adhesion and followed by systems (e.g. Translation, Mitochondria, Lipid synthesis). Individual pathway (row) descriptions are collapsed to respective subsystems as annotated on the right of the heatmap. **(g)** Genes positively correlated with secreted IgG levels across all three high-secreting clones (median *r_s_*>0.1, adjusted *p*<0.05) were identified using bootstrapped correlations (10^2^ iterations ×10^3^-cell subsamples). 15 shared genes were analyzed for protein-protein interaction (PPI) enrichment; non-interacting genes are shown as isolated nodes. Genes previously identified as activated housekeeping genes with IgG production are highlighted as red nodes. **(h)** Genes negatively correlated with secreted IgG (median *r_s_*<−0.1, adjusted *p*<0.05) across all three clones (84 total) form two enriched PPI clusters involving RHO GTPase signaling/trans-Golgi trafficking, and steroid biosynthesis. Genes previously identified as suppressed housekeeping genes with IgG production are highlighted as blue nodes; subcellular localization annotations (endomembrane, cell junction, secreted) are outlined.

To this end, scCellFie metabolic task scoring was combined with protein–protein interaction (PPI) scoring of secretory pathway subprocesses^25,26^ and GO biological process pathway scoring, to quantify biological process activity in each cell. Differentially expressed markers for each high-secreting cluster (adjusted *p* < 0.05, relative to Lo-B/C clones) were further analyzed using Ingenuity Pathway Analysis (IPA) to identify activated or inhibited upstream regulators (|Z-score| >1.5, adjusted *p* < 0.05)^27^. Processes correlated with single-cell IgG secretion (Spearman |*rₛ*| > 0.1, adjusted p < 0.05, relative to Lo-B/C clones) across dominant clusters of all high-secreting clones revealed shared pathways involving translation, amino acid metabolism, lipid metabolism, early-to-late secretory machinery, and adhesion (**Figure 4f, Supplementary Tables 5 and 6**). Krebs Cycle, metabolism of essential (histidine, isoleucine, leucine, phenylalanine, threonine, tryptophan) and non-essential amino acids (alanine/beta-alanine, glutamate, ornithine), and biosynthesis of lipids (glycerophospholipid, cholesterol, sphingolipids) were anti-correlated with IgG secretion, whereas cytoplasmic and mitochondrial translation were positively correlated.

High-secreting clones also displayed suppression of protein folding, ERAD, post-translational modifications, and vesicle trafficking pathways despite elevated translational activity (**Figure 4f**). Additionally, pathway scores for cell adhesion, motility, and cell-cell communication negatively correlated with secretion, consistent with our earlier observation that core protein processing tasks and high-cost secretory proteins are globally suppressed with increasing IgG transcription and secretion (**Figure 3e,h**).

Single-cell gene-level correlation further confirmed these trends: across high-secreting clones, 10 genes were commonly positively correlated with IgG secretion, and 84 genes were commonly anti-correlated (Spearman |*rₛ*| > 0.1, adjusted *p* < 0.05; **Figure 4g,h, Supplementary Table 7**). The top-producing clone (Hi-E) contained 2- to 9-fold greater number of genes significantly correlated with IgG secretion compared to other high producers (Hi-D/F). Positively correlated genes include ribosomal proteins (*Rps15a, Rps3a1*), peptidyl-prolyl cis-trans isomerases catalyzing cis-trans isomerization of proline imidic peptide bonds (*Ppib, Fkbp11*), and a member of the translocon-associated protein (TRAP) family (*Ssr4*). Conversely, shared anti-correlated genes overlapped with suppressed housekeeping genes associated with IgG transcription and secretion (**Figure 3d,e**, blue nodes in **Figure 4h**). These genes were enriched in steroid biosynthesis, RHO GTPase signaling, and trans-Golgi trafficking, and localized to endomembranes, cell junctions, and the secretome—consistent with our previous finding that core machinery and high-cost secPs are suppressed at higher IgG production levels (**Figure 3d-f**). In addition, anti-correlated gene sets from each high-secreting clone significantly overlapped (hypergeometric test, adjusted *p* < 0.05) with PPIs enriched in low rituximab-producing CHO clones^28^ and with genes identified as productivity-enhancing hits in pooled CRISPR KO screens for bispecific antibody production^9,29^ (**Supplementary Figure 6g, Supplementary Table 8**). These results indicate that secretion-coupled transcriptomic trends identified here recapitulate factors previously linked to CHO productivity.

In contrast, low-secreting clones and the parental pool were largely composed of a dominant cluster (cluster 1), enriched for an activated mitotic and reduced translation signature relative to the other clusters (**Supplementary Figure 6a**). Relative to the parental pool, low-secretion clones exhibited lower IgG RNA levels but show activation of mitosis, cell-cell interaction/migration, translation, fatty acid synthesis, ERAD, vesicle trafficking, and oxidative phosphorylation (**Supplementary Figure 6c-e**). The low-secreting clones thus show a proliferative and stress-associated (e.g. elevated ERAD/protein processing) phenotype compared to the high-secreting clones and to the parental pool.

Overall, the shared anti-correlation of multiple metabolic and secretory task scores with single-cell IgG secretion, together with common activated and inhibited regulators, suggests that high-secreting clones share a translation-associated expression program linked to IgG secretion. In contrast, low-secreting clones show comparatively higher activity signatures for mitosis/cell-cell interactions and multiple protein-processing pathways.

### Clonal variation among high-secreting clones reveals distinct metabolic strategies underlying translation-focused phenotypes

High secreting clones shared a consensus of suppressed energy and protein processing-associated tasks supporting a translation-optimized phenotype. However, variation in metabolic task activity relative to low clones (**Figure 4d**, **Figure 5b**), and distinct composition of transcriptomic cells states (**Figure 2j**), indicate that multiple regulatory routes can support efficient IgG production. Furthermore, despite all clones being predominantly composed of G1-phase cells (cell-cycle phase regressed, **Supplementary Figure 4b,d-e**), divergence in marker genes and enriched pathways among the 9 transcriptomic clusters (**Supplementary Figure 6a,b, Supplementary Table 9**) support functionally distinct cell states during the highly productive phase within the fed-batch culture.

**Figure 5.**
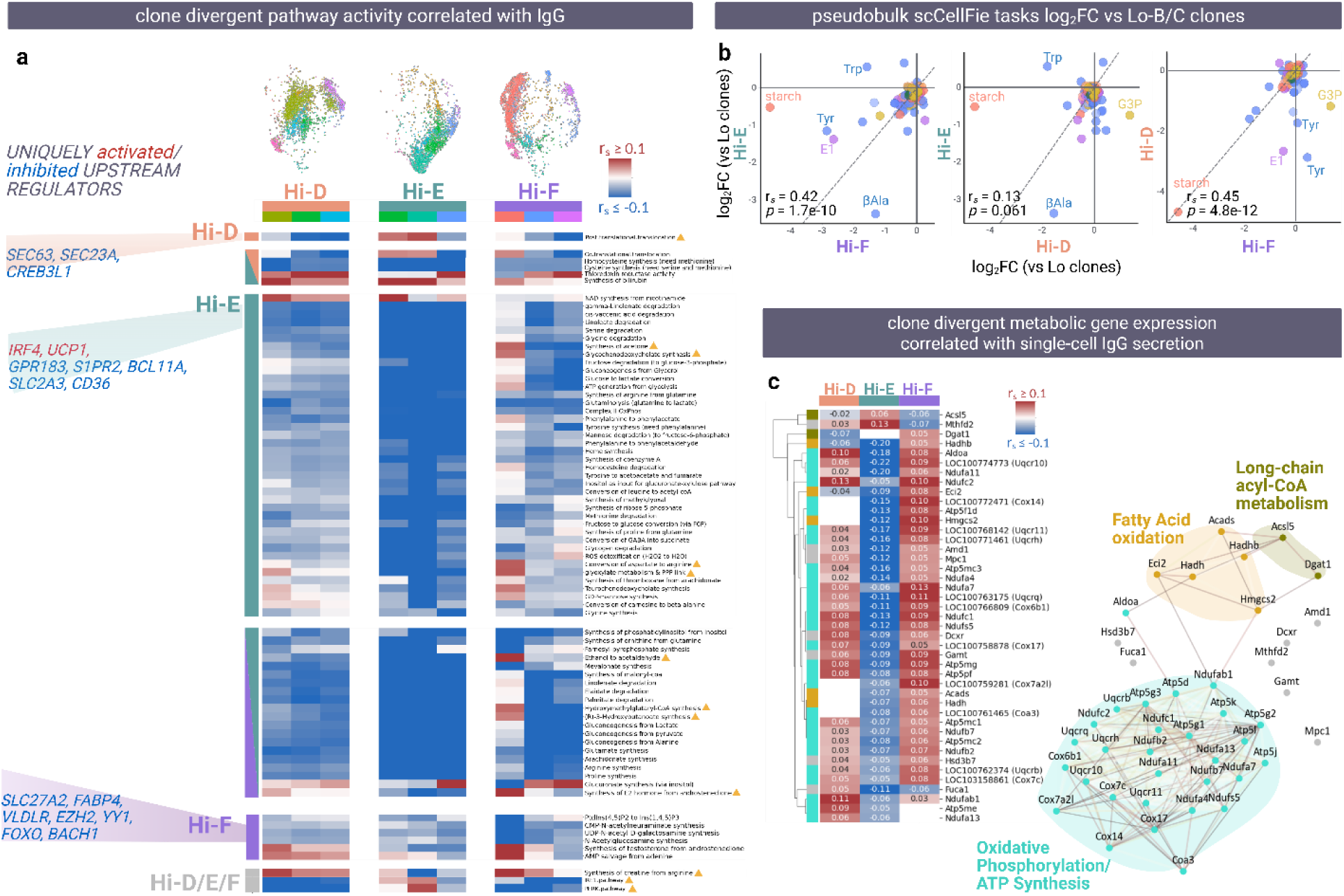
High-secreting clones diverge in energy and redox metabolic phenotypes coupled with single-cell IgG secretion. (**a**) Heatmap of pathway activities significantly coupled with IgG secretion (Spearman correlation, |*r_s_*| > 0.1, adjusted *p* < 0.05, in at least 1 top cluster relative to the low-secreting clones) and uniquely present in 1 or 2 high-secreting clones. Pathways are grouped by clone-specific behavior, including cases of opposite correlation directionality as highlighted by the yellow triangles. Grey bar indicates significant pathway activity-IgG coupling in all 3 clones (Hi-D/E/F). Clone-specific enriched upstream regulators from IPA are annotated. **(b)** Pairwise Spearman correlations of log₂FC in pseudobulked scCellFie metabolic task scores for each high-secreting clone (Hi-D/E/F) relative to the low-secreting clones (Lo-B/C). Each panel shows a pairwise comparison between two high-secreting clones; the dotted line denotes the *y = x* reference. All significance values reflect Benjamini–Hochberg–adjusted p-values. Top differentially expressed metabolic subsystems are annotated. (Tyr: tyrosine metabolism, Trp: tryptophan metabolism, βAla: beta-alanine metabolism, E1: E1 estrogen metabolism, G3P: glycerol-3-phosphate metabolism) **(c)** Heatmap of metabolic genes annotated under significantly IgG-coupled scCellFie tasks and showing clone-specific coupling directionality (median bootstrapped Spearman rho |*r_s_*| > 0.1, adjusted *p* < 0.05, relative to the low secreting clone populations). 43 genes were evaluated for PPI enrichment, with enriched pathway clusters highlighted as node colors, background shade, and heatmap row colors (grey: nodes outside of enriched cluster pathways).

To unravel how these clone-specific cell states relate to IgG secretion, we performed cluster-resolved correlation analysis of metabolic and secretory pathway activities with single-cell IgG levels for the top 3 clusters per high-secreting clone (Spearman |*r_s_* | > 0.1, adjusted *p* < 0.05, relative to Lo-B/C clones, **Figure 5a, Supplementary Table 10**). Consistent with the consensus analysis, clone Hi-E exhibited the strongest anti-correlation between IgG secretion and amino acid (glutamate, glycine, methionine, serine), lipid (fatty acid, eicosanoid, cholesterol), and carbohydrate-to-ATP metabolic tasks, alongside positive coupling with IRE1/PERK signaling, co-and post-translational translocation, NAD synthesis from nicotinamide, and bilirubin synthesis. In contrast, clone Hi-F uniquely activated carbohydrate utilization pathways, including glycolysis/gluconeogenesis and the glyoxylate–pentose phosphate axis. Clone Hi-F also showed increases in AMP salvage and lipid-metabolic reactions including HMG-CoA and glycochenodeoxycholate synthesis. Clone Hi-D displayed negative coupling of IRE1/PERK activity with IgG secretion (opposite to Hi-E), shared elevated NAD salvage with Hi-E, and showed more modest coupling between carbohydrate metabolism and ATP synthesis compared to Hi-F (**Figure 5a**).

To dissect the gene-level basis of these differences, we examined expression of scCellFie genes involved in significantly IgG-coupled metabolic tasks. 43 genes displayed a significant correlation with IgG secretion in one high-secreting clone but the opposite correlation direction in at least one other clone (**Figure 5c, Supplementary Table 11**). Among these clone-unique couplings, *Acsl5* and *Mthfd2* showed positive correlations with IgG secretion specifically in Hi-E, but were anti-correlated in Hi-D and Hi-F (**Figure 5c**). *Acsl5* catalyzes conversion of long-chain fatty acids to acyl-CoA intermediates that can be directed toward complex lipid biosynthesis, membrane remodeling, or mitochondrial β-oxidation^30,31^, and *Mthfd2* encodes a mitochondrial one-carbon enzyme linked to redox and nucleotide metabolism^32–34^. Clone Hi-E displayed a predominantly suppressed lipid synthesis signature but positive coupling of *Acsl5* and IgG secretion, along with positive coupling of IgG secretion with *Mthfd2* and NAD salvage (NAD synthesis from nicotinamide) with IgG secretion (**Figure 5a,c**). This transcriptomic signature coincided with a lower growth rate, yet the highest IgG transcript and secretion levels across clones (**Figure 2c–e**, **Figure 5b**).

We next asked how the broader set of 43 clone-divergent genes organized at the protein-interaction level. STRING protein-protein interaction network analysis revealed three main clusters: (i) oxidative phosphorylation/ATP synthesis, (ii) fatty-acid β-oxidation, and (iii) long-chain acyl-CoA metabolism (**Figure 5c**). Within this network, *Aldoa, Ndufab1, Hmgcs2, Hadh*, and *Eci2* occupied central positions connecting oxidative phosphorylation genes with those involved in mitochondrial fatty-acid metabolism (**Figure 5d**). In clone Hi-E, expression of these central nodes was negatively correlated with IgG secretion, whereas clones Hi-D and Hi-F showed moderate to strong positive coupling of these nodes with IgG secretion **Figure 5c,d**). Consistent with these clone-specific coupling patterns, clone Hi-F—which exhibited the highest growth rate (**Figure 4a**)—showed unique positive correlations between IgG secretion and respiratory chain subunits (*LOC100772471/Cox14, LOC100759281/Cox7a2l, Atp5f1d*) as well as multiple β-oxidation genes (*Hmgcs2, Gamt, Eci2, Acads, Hadh, Hadhb*) (**Figure 5a,c**). In contrast, clone Hi-D exhibited lower IRE1/PERK coupling and weaker correlations between IgG secretion and oxidative phosphorylation/mitochondrial lipid metabolism module genes relative to Hi-E and Hi-F, together with an intermediate growth profile (**Figure 4a**,**Figure 5a,c**).

Together, SEC-seq combined with targeted metabolic–secretory pathway analysis revealed cluster-resolved, IgG-coupled clonal variation among high-secreting clones. This analysis identified clone-specific pathway–secretion associations and gene sets that organized into energy metabolism modules (oxidative phosphorylation and mitochondrial lipid metabolism), identifying candidate metabolic and secretory nodes that differ across clones.

## Discussion

Single-cell omics is transforming our understanding of CHO genotype–phenotype relationships, but most work has focused on transcriptomic heterogeneity, clonal stability, or pathways associated with bulk phenotypes and recombinant transcription rather than recombinant protein secretion. Here, we used SEC-seq to link the transcriptome to recombinant IgG secretion at single-cell resolution in CHO cells. To our knowledge, this is the first published application of SEC-seq in CHO clones, enabling direct interrogation of transcriptional, metabolic, and secretory programs coupled with IgG secretion in a biomanufacturing-relevant context.

Applying SEC-seq to a parental pool and clones spanning a range of bulk cell-specific productivities, we observed that the high-secreting clones each showed distinct transcriptomic clusters, whereas the low-secreting clones and the parental pool showed more similar cluster compositions. Across all cell lines, IgG heavy- and light-chain transcripts accounted for up to ∼1-40% of the single-cell transcriptome, imposing substantial transcriptional and secretory burden, consistent with prior transcriptomic studies in recombinant CHO cells^9,21,22^ and IgG-secreting plasma cells^20^. IgG transcription and secretion were coupled primarily at the population and clonal levels but weakly at the single-cell or subpopulation level within a given clone. This decoupling likely reflects differences in mRNA and protein turnover dynamics^35–38^. While IgG transcript abundance is coupled with overall productivity, it did not strongly discriminate transcriptomic subpopulations. This motivated an investigation of metabolic and secretory processes adapted to handle recombinant burden, focusing on signatures shared among high producers as potential engineering targets and clone-specific signatures as distinct “paths to success,” which may serve as more informative markers of cellular “readiness” for high productivity.

We first assessed how core cellular machinery adapts to the demands of IgG production by correlating housekeeping gene expression with IgG transcript and secretion across clones. Globally, housekeeping expression remained stable between low- and high-IgG secreting subpopulations, indicating preservation of essential functions under high recombinant load. However, a subset of housekeeping genes showed coordinated shifts across low- and high-secreting cells: translation and protein-targeting enriched pathways were activated with IgG, whereas vesicle trafficking, secretion, and high-ATP-cost endogenous secretory proteins were suppressed. This likely reflects selective downregulation of non-essential endogenous secretion—rather than impaired recombinant secretion—thereby prioritizing IgG synthesis and export. Similar rebalancing has been reported in bulk analyses of high-producing CHO and hybridoma lines and in highly secretory human tissues, suggesting a conserved adaptive in mammalian cells^21,22,24^.

Extending beyond housekeeping genes, we asked how genome-wide expression impacts single-cell IgG secretion, and how high-secreting clones achieve further productivity gains. Across high-secreting clones, genes significantly anti-correlated with IgG secretion outnumbered positively correlated genes, consistent with prior findings that many productivity-associated markers are downregulated in high producers^21,22^. To assess whether these anti-correlated genes converge on previously reported productivity associated markers and functional perturbation hits, we compared our clone-specific and shared anti-correlated gene sets to published interactome enrichment in high vs. low rituximab-producing CHO clones^28^ and CRISPR KO productivity screens in bispecific antibody-producing CHO cells^9,29^ (**Supplementary Figure 6g**). PPIs enriched from low producers and gene knockouts leading to increased bsAb productivity significantly overlapped with our anti-correlated markers, particularly in pathways related to cell motility, long-chain fatty-acid metabolism, ER-associated degradation (ERAD), and vesicle trafficking. These results show that SEC-seq can resolve and extend candidate negative regulators identified by functional screens, and support efforts toward “minimal” CHO genomes in which non-essential protein-processing tasks are removed to increase productive capacity^39–42^.

We next contextualized these gene-level trends at the pathway level by relating metabolic and secretory task activities to single-cell IgG secretion. Across all three high-secreting clones, we observed extensive suppression of metabolic and secretory tasks that were significantly anti-correlated with IgG secretion relative to low producers. High clones converged on a translation-optimized state during the productive phase, marked by increased cytoplasmic and mitochondrial translation. This state was accompanied by suppression of lipid biosynthesis, vesicle trafficking, adhesion, and cell-junction pathways which are likely less essential in suspension-adapted CHO cells and may reallocate ATP towards for IgG production. Early-to-late secretory pathway activity, including proteostasis and post-translational modifications processes, was also reduced despite elevated translation, consistent with efficient proteostasis and lipid homeostasis without sustained ER stress. These trends align with prior observations that high-producing CHO cells downregulate energetically expensive endogenous secreted proteins, including GPI-anchored proteins, to prioritize recombinant secretion^24,43^. Conversely, we found that low-secreting clones remain growth- and stress-focused, likely exhibiting reduced capacity to suppress unnecessary pathways and to efficiently funnel resources toward IgG production.

Despite this convergent translation-focused program, high-secreting clones showed clonal variation in transcriptomic cluster composition and metabolic task activity, indicating different regulatory routes to similarly high productivity. Clone Hi-E, which had the highest IgG transcription and secretion and the lowest growth rate, showed pronounced anti-correlation between IgG and amino-acid, lipid, and carbohydrate-to-ATP metabolic tasks, alongside positive coupling of IRE1/PERK signaling, co- and post-translational translocation, and NAD salvage. In contrast, clone Hi-F, with the highest growth rate, showed strong positive coupling between IgG and glycolysis/gluconeogenesis, the glyoxylate–pentose phosphate axis, respiratory chain components, and mitochondrial β-oxidation, consistent with an oxidative, fatty acid–driven energy strategy. Clone Hi-D exhibited an intermediate phenotype, with opposite IRE1/PERK coupling relative to Hi-E, elevated NAD salvage, and more modest coupling of central carbon metabolism to IgG than Hi-F.

To connect these clone-level pathway patterns to specific molecular drivers, we identified 43 metabolic genes whose IgG–expression correlations differed across high clones and enriched for protein–protein interaction modules under oxidative phosphorylation/ATP synthesis, fatty-acid β-oxidation, and long-chain acyl-CoA metabolism. Within this network, *Aldoa*, *Ndufab1*, *Hmgcs2*, *Hadh*, and *Eci2* occupied central positions linking glycolysis^44^, mitochondrial respiration^45–47^, and fatty-acid metabolism^45–50^. These genes showed negative coupling to IgG in Hi-E but moderate-to-strong positive coupling in Hi-D/Hi-F, revealing a molecular “bridge” mirroring the clone-level differences in mitochondrial energy strategies. The positive coupling of respiratory chain and β-oxidation genes with IgG in Hi-F is consistent with CHO flux and modeling studies linking increased TCA/oxidative phosphorylation (often with elevated pentose phosphate activity) to high antibody productivity and q ^51,52^. Conversely, the inverse coupling observed in Hi-E suggests that this clone achieves high IgG output through a low-growth, mitochondria-attenuated strategy, rather than enhanced oxidative metabolism.

Low-growth, high-specific-productivity phenotypes have been characterized in CHO cells and support a rebalancing of cellular resources from biomass generation toward translation and secretion ^53–55^. We therefore examined genes uniquely positively coupled to IgG in Hi-E, identifying *Acsl5* and *Mthfd2* as potential clone- and phenotype-specific metabolic levers. *Acsl5*—a long-chain acyl-CoA synthetase localized to ER and mitochondria—showed positive coupling with IgG secretion despite broader suppression of lipid synthesis signatures, suggesting preferential channeling of acyl-CoA toward lipid pools that support ER remodeling and vesicle trafficking rather than bulk membrane expansion for biomass^30,56–58^. *Mthfd2*, a mitochondrial one-carbon enzyme, was also uniquely positively coupled to IgG in Hi-E. Although its role in CHO secretion is not established, *Mthfd2*—involved in one-carbon and redox metabolism—is upregulated across many cancers^34^, and its inhibition disrupts redox homeostasis, glycolysis, mitochondrial respiration, and UPR signaling under highly secretory cancer models (i.e. multiple myeloma)^32,33^. A similar NAD/one-carbon axis in Hi-E may supply ATP, NAD(P)H, and formate to sustain high translation under low growth, highlighting a potential metabolic marker for follow-up perturbation in CHO. Together, these patterns show that high-secreting clones share a translation-focused, stress-attenuated program relative to low producers but differ in how they balance glycolysis, oxidative phosphorylation, fatty-acid oxidation, and UPR signaling to support secretion.

The observation that clone Hi-E shows both the tightest transcriptional coupling with IgG secretion and the highest number of anti-correlated genes is consistent with minimal-genome models^39,41,42^ in which non-essential functions are attenuated and a larger fraction of the cellular “budget” is dedicated to translation and secretion. An open question, however, is whether these suppressed genes and pathways are truly dispensable under production conditions, or whether some are directly or indirectly required to achieve or maintain high productivity and optimized to lower expression. These questions are well suited to systematic perturbation studies using CRISPRi/KO screens^9,29,59,60^ and genome-scale modeling^24,61,62^ to further probe metabolic nodes such as *Aldoa, Ndufab1, Acsl5,* or *Mthfd2* in clone-specific contexts.

This study has limitations that motivate future work. SEC-seq was applied at a single time point during peak production in a dynamic fed-batch process, providing only a snapshot of clonal and subpopulation states. Given the genomic and epigenetic plasticity of CHO cells^6,63^, it will be important to assess how stable these transcriptional and metabolic programs are across process phases or long-term culture. As the field increasingly adopts transposase- or landing pad–mediated targeted integration to reduce clonal heterogeneity, it also remains unclear whether the diversity in transcriptomic and metabolic wiring observed here will persist or become constrained. Longitudinal multi-omics profiling across process stages^12,55,64–66^, coupled with controlled integration strategies^67^, can help address these questions.

Nevertheless, our work illustrates how SEC-seq can bridge transcriptional states and secretion phenotype at single-cell resolution, and, together with pathway activity modeling and network analysis, identify shared and clone-specific signatures by which CHO cells accommodate recombinant IgG burden. We anticipate that integrating methods like SEC-seq with high-throughput CRISPR perturbation and rational chassis engineering can move the field toward “precision wiring” of CHO cell factories that robustly deliver desired combinations of growth, productivity, and product quality for diverse biologics.

## Materials and Methods

### CHO cell line generation and passaging

Cell lines were generated by transfecting a proprietary CHO-K1 glutamine synthetase knockout host with plasmid DNA encoding a model human IgG1 antibody using standard cell line development procedures^17,68,69^. After transfection, a stably expressing pool was established by repeated passaging in proprietary selective medium lacking glutamine. Single-cell derived clones were isolated from this pool using the Beacon Instrument^17,68,69^ (Bruker Cellular Analysis, Inc., Emeryville, CA) and expanded and cultured in 96-well plates (Corning, NY), 24-deep well plates (Corning, Corning, NY), or 50 ml spin tubes (TPP, Trasadingen, Switzerland) in growth media under standard culture conditions (36°C, 5% CO_2_ and 85% humidity). Cell banks were created by freezing 1 × 10^7^ viable cells in growth medium with 10% DMSO (Sigma-Aldrich, St. Louis, MO). Cells were maintained by passaging every 3 or 4 days.

### Fed-batch Production Assays

Cell lines were evaluated in small-scale fed-batch cultures in 50 ml spin tubes (TPP, Trasadingen, Switzerland) with a 20-30 mL working volume. Cells were incubated at 36°C, 5% CO_2_, 85% relative humidity and shaken at 225 rpm (50 mm orbital diameter, Kuhner ISF4-X incubator, Kuhner AG, Basel, Switzerland). Proprietary production and feed media were used, and culture received bolus feeds on days 3, 6, and 8. No active pH and dissolved oxygen control were applied. In-process samples were collected on days 0, 3, 6, and 8 for fed-batch experiments supporting SEC-seq analysis or daily for spent media analysis. Viable cell density and viability were determined using a Vi-Cell Blu cell counter (Beckman Coulter, Brea, CA). IgG titers were quantified by affinity Protein A high performance liquid chromatography (HPLC) (Protein A, Waters, Milford, MA) using cell culture supernatant. Spent media routine metabolites were analyzed using the BioProfile FLEX2 analyzer (Nova Biomedical, Waltham, MA) and amino acids were measured by REBEL CE-MS (Repligen, Waltham, MA).

### Nanovial loading and functionalization for SEC-seq

#### Reagents and Nanovial Preparation

35µm biotinylated gelatin nanovials (Partillion Bioscience, Pasadena, CA) were used for all SEC-seq method development and experiments according to manufacturer guidelines and protocols based on adherent-cell secretion applications^19,20,70^. Nanovial functionalization for SEC-seq IgG capture incorporated a 1:10 ratio of oligo-barcoded streptavidin (Biolegend TotalSeq™-B0975 #405369, San Diego, CA; Thermo Fisher PN#434302, Waltham, MA) to unbarcoded streptavidin (Thermo Fisher #434302, Waltham, MA), biotinylated anti-IgG capture antibody (Southern Biotech #2014-08, Birmingham, AL), and an oligo-barcoded anti-PE detection antibody (BioLegend TotalSeq-B0911 #408113 San Diego, CA). Proxy reagents of streptavidin with no oligo-barcode and APC anti-PE (Biolegend TotalSeq™-B0911 #408108 San Diego, CA) in place of oligo-barcoded anti-PE detection antibody were used during optimization. Cytometry profiles were acquired on a MACSQuant Analyzer 10 (Miltenyi Biotec, Bergisch Gladbach, Germany) and analyzed in FlowJo (FlowJo, LLC, Ashland, Oregon). Final reagent concentrations of oligo-barcoded streptavidin, streptavidin, and capture antibody aligned with Partillion Bioscience protocols were selected to saturate biotinylated nanovials.

#### Cell Loading into Nanovials

CHO cells washed with Ca²⁺/Mg²⁺ DPBS + 0.4% BSA were resuspended in fresh growth media at 1.5 x 10^6^ cells/mL and pipette mixed with nanovials at a 4:1 to 5:1 final cell-to-nanovial ratio. Suspensions were incubated at 36°C, 5% CO2 for 1 hour in a static incubator to allow cells adherence within nanovial cavities, then washed and strained using CellTrics 20µm strainers (Sysmex Partec GmbH #040042325, Gorlitz, Germany) according to Partillion Bioscience manufacturer protocols following Partillion Bioscience protocols. Calcein AM cell-permeant dye (Thermo Fisher #C3100MP, Waltham, MA) was added to target 0.75 µM effective concentration about 30 minutes prior to imaging or sorting for GFP-positive events. Brightfield and GFP images of cell-loaded nanovials were acquired using an EVOS microscope (Thermo Fisher, Waltham, MA).

#### Detection Antibody Optimization

Streptavidin/anti-IgG capture-antibody–functionalized nanovials were incubated with serially diluted crude harvests (1–80 ng/mL IgG, containing concentrations below and above expected bulk cell specific productivities for typical CHO clones during the production phase of fed-batch culture) for 1 h at 36°C, followed by PE–anti-IgG staining for 0.5 h (80–160 µg/mL range tested). The crude harvest initial stock was quantified using affinity Protein A high performance liquid chromatography (HPLC) (Protein A, Waters, Milford, MA). PE median fluorescence intensity (MFI) was correlated with theoretical IgG using Spearman’s rho to define the quantifiable dynamic range. Final optimized concentrations were 120 µg/mL (PE anti-IgG) and 130 µg/mL (anti-PE). Validation was performed using wild-type, low-secreting, and high-secreting clones (Days 7–9) to confirm MFI–q_p_ concordance (*r_s_* > 0.9). Final antibody concentrations were validated using cell-loaded and functionalized nanovials using cells from non-expressing wild-type host cell line, low-secreting clones, and high-secreting clones sampled during Day 7-9 of a fed-batch culture to assess Spearman correlation (*r*_s_ > 0.9) of MFI with bulk cell specific productivity. Similar experiments and validation were conducted for allophycocyanin (APC) anti-PE detection antibody as a proxy for oligo-barcoded anti-PE antibody. Optimization of detection antibodies resulted in 120 µg/mL and 130 µg/mL effective concentrations for PE anti-IgG antibody and anti-PE antibodies, respectively.

### SEC-seq application to CHO cells

The optimized SEC-seq workflow was applied to 7 cell lines—non-producing wild-type host, parental pool cell line (Pool-A), 2 low secreting clones (Lo-B/C), and 3 high secretion clones (Hi-D/E/F)—on Day 8 of fed-batch production. For each sample, 6 x 10^6^ nanovials were mixed and incubated with 30 x 10^6^ cells. An aliquot of Pool-A cell suspension (free cells) was retained in 4°C as a control for 10X Genomics processing.

Cell-loaded nanovials were strained, functionalized, and incubated for IgG secretion capture. After detection antibody staining, cell-loaded nanovials were sorted (GFP⁺/FSC⁺ gate) on a BD FACSFusion (100-µm nozzle) calibrated for drop delay according to Partillion Bioscience guidance protocols. Sorted nanovials were concentrated to target 500 nanovials/µL and loaded into 10x Genomics Chromium Chip G (16.5–30 µL) to target 5,000–8,000 cells. The loading procedure was modified to account for nanovials settling more quickly than cells^19^. Free-cell controls were loaded per 10X Genomics manufacturer protocols.

Gene expression (GEX) and feature-barcode (PRO) libraries were generated using the Chromium Next GEM Single Cell 3’ v3.1 Dual-Index Kit with Feature Barcoding (10X Genomics, Pleasanton, CA). Libraries were QC-assessed using TapeStation4200 (Agilent Technologies, Santa Clara, CA), quantified using Qubit Flex (Thermo Fisher, Waltham, MA), pooled at a 5:1 GEX:PRO ratio, spiked with 1% PhiX, and sequenced (2 × 150 bp) on a NextSeq2000 using P4 XLEAP-SBS flow cell (Illumina, San Diego, CA).

### Analysis of SEC-seq data

#### Alignment and Preprocessing

GEX and PRO FASTQ files were processed with CellRanger v6.1.2 *count* considering streptavidin and detection antibody oligo barcodes. These were aligned to CriGri-PICRH-1.0 reference^71^ extended with CHO-K1 mitochondrial genome^72^ (GenBank: KX576660) and proprietary IgG plasmid sequence prepared using CellRanger *mkref*. Sample count matrices were aggregated without normalization using CellRanger *aggr* and imported into Seurat^73,74^ (v4.3.0). Cells were classified as nanovial-loaded if >10³ streptavidin oligo counts were detected. PRO counts were CLR-normalized^75^. Cells were filtered by: log_10_ genes per UMI > 0.78, nCount (number of UMIs or transcripts) > 500, nFeature_RNA (number of genes) > 1500, mitochondrial fraction < 0.6. Genes were kept if ≥10 cells contained non-zero counts.

Cell-cycle scoring and regression were performed with *CellCycleScoring* and SCTransform in Seurat. Expression was normalized using SCTransform. IgG HC/LC genes were excluded from dimensionality reduction to avoid bias in UMAP projection and cluster generation. UMAPs and Louvain clustering were generated using default Seurat workflows.

#### Single-cell pathway scoring, pathway- and gene-level single-cell correlations

Pathway activity scores were computed using Seurat’s *FeatureExpression* and scCellFie^76^, and protein-protein interaction (PPI) scores were computed using ORIGINS2^25^ with SCTransform-normalized data. Gene sets were obtained from GO-BP^77^ and secRecon^22^. scCellFie metabolic task scoring used human gene-protein-reaction (GPR) mappings^43,76,78^. secRecon PPI networks were built from PCNet^79^.

For initial scCellFie group-level comparisons, pseudobulk profiles were generated by averaging task scores over non-overlapping batches of 20 randomly sampled cells within each sample–cluster group (≥20 cells required). Pseudobulk matrices were used for PCA and differential pathway activity testing. Group comparisons (e.g., high- vs low-secreting clones) were performed using Welch’s t-tests on pseudobulk values, with log₂ fold changes computed from group means. *p*-values were corrected using the Benjamini–Hochberg FDR procedure.

For gene–IgG correlations, each high-secreting clone (Hi-D/E/F) was compared to an equal number of cells sampled from the combined low clones (Lo-B/C). To address cell count imbalance, nonparametric bootstrapping (100 iterations × 1,000 cells or max available) was applied. In each bootstrapped dataset, Spearman correlations (*r*_s_) were computed and aggregated using median *r*_s_ and Fisher-combined *p*-values with Benjamini–Hochberg correction. Overrepresentation of published CHO genesets with Hi-D/E/F positively and negatively correlated genes with IgG (|*r*_s_ | > 0.05, BH *p* < 0.05) were conducted using hypergeometric test. Published genesets were results from FcBAR proteome, transcriptome, and regression analysis (filtered for adjusted *p* < 0.05) of high and low rituximab producing CHO clones (Wu et al.^28^), enriched CRISPR gene knockouts (KOs) enhancing productivity (>5% increase in normalized titer) for 2 bispecific antibody (bsAb) producing CHO cell lines (Bauer et al.^9^), and significant CRISPR KOs enriched for bsAb producing CHO cells (Kim et al.^29^). Pathway enrichment for overlapping genes was performed using gseapy *enrichr* and filtered for adjusted *p* < 0.05 significant KEGG and GO terms.

Pathway scores were similarly collated to perform Spearman correlation against single-cell IgG protein levels. Differential expression for clusters and pairwise comparisons were assessed using Seurat *FindAllMarkers*/*FindMarkers* with Wilcoxon tests (adjusted *p* < 0.05). Enrichment analyses were performed using clusterProfiler (GO-BP) and IPA (Qiagen, Germantown MD). STRING (*mus musculus*) was used for PPI enrichment^80,81^. Activated and inhibited upstream regulators were identified with IPA (|Z| ≥ 1.5, adjusted *p* < 0.05).

Consensus “high-clone” pathways were defined as those with significant single-cell pathway-IgG correlations (|*r_s_*| > 0.1, adjusted *p <* 0.05) in ≥1 of the top three abundant clusters of every high-secreting clone. Clone-unique pathways were significant in ≥1 cluster of that clone only or exhibiting opposite coupling directionality from the other clones. Visualizations (density/scatter plots, heatmaps, Venn diagrams, UpSet plots, network graphs) were performed in Python (seaborn, matplotlib, upsetplot, NetworkX) and R (Seurat, ComplexHeatmap, ggplot2).

### Housekeeping gene and secreted protein analysis

CHO housekeeping genes were previously identified by Joshi and colleagues^23,82^. Gene-level Gini coefficients (*G*) were computed using the GiniWegNeg R package based on counts-per-million (CPM) expression and SCTransform-normalized data. Significance of reduced variability among housekeeping genes relative to randomly sampled genes (size-matched) was assessed using 5,000-iteration bootstrap tests and 10,000-permutation label-reshuffling tests. Because curated housekeeping lists can include genes that are not uniformly expressed across experimental conditions and CHO hosts, we defined a conservative low-variability subset (“Gini genes”) as housekeeping genes whose Gini coefficients fell within the lowest 20th percentile of the bootstrapped global (random gene group) Gini distribution^23^ and used this subset for downstream analysis.

High- (q90) and low- (q10) IgG-secreting subpopulations were identified for each cell line group (pool, low-secreting clones, high-secreting clones) based on the 90^th^ and 10^th^ percentile of SEC-seq IgG CLR-transformed counts per group, respectively. Gini coefficients for these groups were compared using one-way ANOVA.

Spearman partial correlations (controlling for sequencing depth for LC/HC RNA) were computed between housekeeping gene expression and IgG LC RNA, HC RNA, or secreted IgG abundance. Gini genes with *r_s_* < 0 (adjusted *p* < 0.05) were classified as suppressed with increasing IgG levels, whereas genes with *r_s_* > 0 were classified as activated with increasing IgG levels. GO-BP enrichment of activated/suppressed genesets was performed using clusterProfiler against background geneset for all genes in the initial counts matrix post-QC/filtering. GO-BP enriched terms were manually collapsed into major biological pathway themes and visualized using seaborn v0.11.2.

Secreted protein genes by CHO (secP) and ATP costs previously calculated by Gutierrez and colleagues^24^ were mapped to CriGri-PICRH-1.0 gene symbols. Gene lengths were extracted from CriGri-PICRH-1.0 GTF annotation file.

### Spent media analysis

Daily metabolite (glucose, glutamine, glutamate, lactate, ammonium; amino acids) and amino acid measurements were modeled using phase-resolved ODE fitting across early exponential (Day 0–3), late exponential (Day 3–6), and production (Day 6–10) phases^55^.

Extracellular metabolite concentrations were modeled using a volume-explicit mass balance:

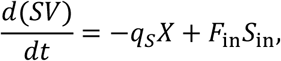

where 𝑆 is the extracellular metabolite concentration, 𝑉 is culture volume, 𝑋 is the total viable biomass, 𝑞_𝑆_ is the cell-specific consumption/production rate, 𝐹_in_ is feed rate, and 𝑆_in_ is the metabolite concentration in the feed. Volume dynamics followed:

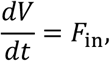

and viable cell growth was described by:

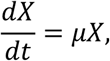

with phase-specific growth rates 𝜇. Parameters 𝑞_𝑆_ and 𝜇 were fit separately for each growth phase and clone using nonlinear least squares. Clone-level differences metabolite (glucose, glutamine, glutamate, lactate, ammonium) and amino acid fitted production phase consumption rates were assessed with principal component analysis.

## Acknowledgements

The authors thank colleagues in Amgen Drug Substance Technologies, Attribute Sciences, Discovery Biological Sciences, and Cytometry Sciences for their invaluable project support, especially Kristi Daris, Natalia Gomez, Eugenia Zah, Maria Cuellar, Pheng Yam, Jonathan Diep, Anne Abrajano, Charilyn Tejamo, Kayley Cox, Becky Cheng, Ewelina Zasadzinska, John Ferbas, Lukasz Ochyl, Andres Lopez, Asha Kelley, and the Rapid Analytics team. The authors also thank Joseph De Rutte, Jesse Liang, Katie Kershaw, and Shreya Udani from Partillion Bioscience for providing technical support for SEC-seq protocols and development.

This work was supported by funding generously provided by NIGMS (R35 GM119850) and Amgen.

## Author Contributions

J.T., F.L., and N.E.L. contributed to the experimental design. J.T. generated the cell lines, performed the experiments, and conducted the data analysis. J.T wrote the original draft. J.S. and N.E.L. provided project support and advising. All authors provided critical feedback and helped shape the research, analysis, and manuscript.

## Lead Contact

Further information and requests for resources and reagents should be directed to and will be fulfilled by the Lead Contact (Nathan E. Lewis).

## Materials availability

This study did not generate new materials.

## Data and Code availability

Data is provided in the supplementary material This paper does not report original code.

## Declaration of generative AI and AI-assisted technologies in the writing process

During the preparation of this work the authors used ChatGPT5 in a limited manner to improve the readability and language of the manuscript. After using this tool, the authors reviewed and edited the content. The authors take full responsibility for the content of the published article.

---

**Supplementary Figure 1.**
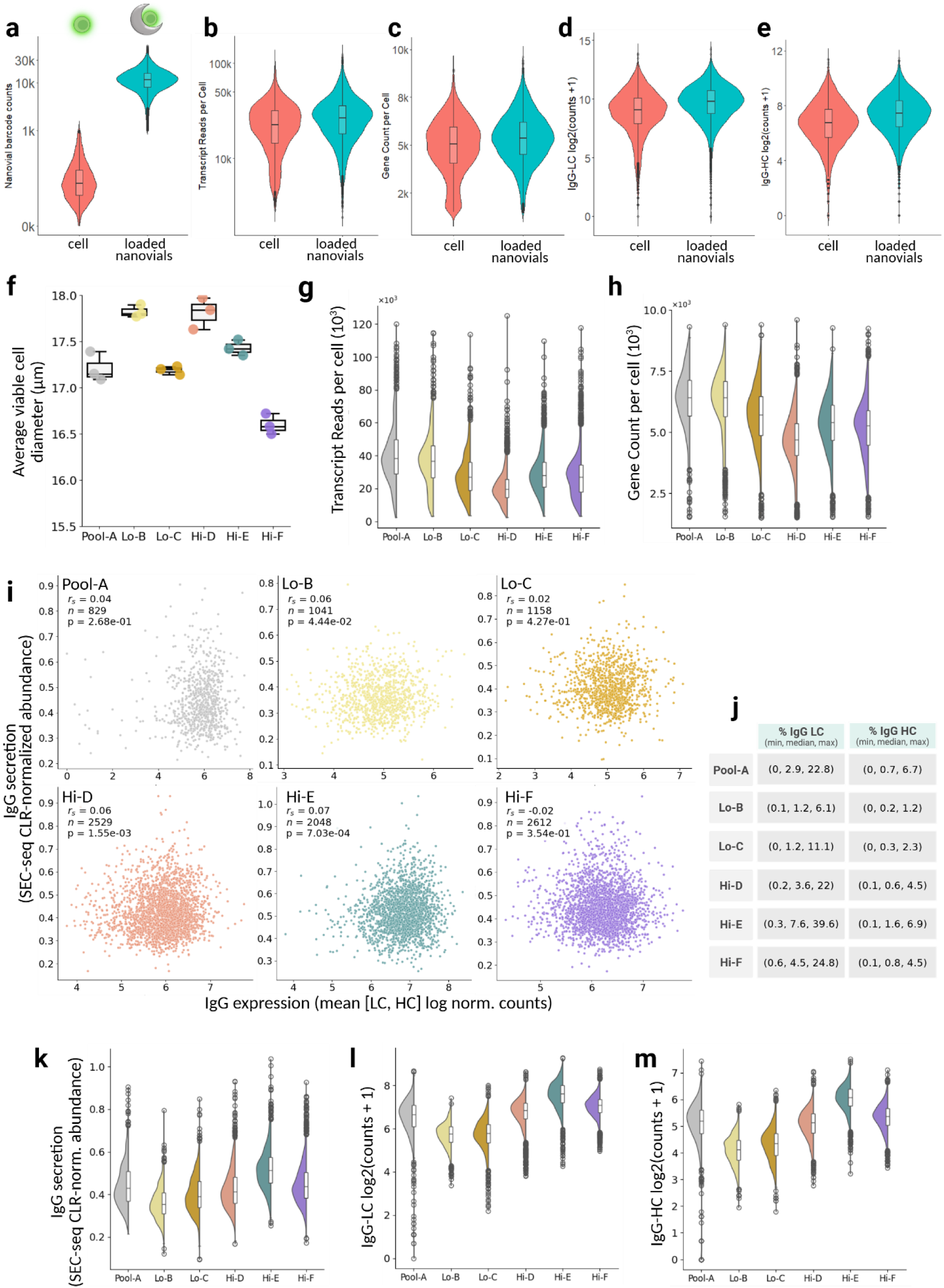
10X Genomics single-cell transcriptome and feature barcoding libraries generated from cells in nanovials achieving adequate sequencing depth and quality. (**a-e, top**) Comparison of free cells and cell-loaded nanovials across sequencing QC metrics (5 clones + 1 parental pool). Metrics shown include: (**a**) streptavidin feature-barcode counts distinguishing free versus loaded cells; (**b**) number of individual transcripts or UMIs (unique molecular identifiers) per cell; (c) number of genes detected per cell; (**d**) IgG light-chain (LC) transcript levels per cell, and (**e**) IgG heavy-chain (HC) transcripts. (**f**) Average viable cell diameter (N=3) for each cell line sample on the day of SEC-seq workflow (Day 8 of small-scale fed-batch assay). **(g–m, bottom)** Post-filtering QC metrics for SEC-seq (cell-loaded nanovials) transcriptome and feature barcode libraries across samples: (**g**) number of individual transcripts or UMIs (unique molecular identifiers) per cell, (h) number of genes detected per cell, (**i**) scatterplots of single-cell IgG expression (mean of IgG LC and HC expression) versus single-cell IgG secretion with Spearman correlation coefficient (*r_s_*), number of cells (*n*), and p-values (*p*) annotated. All significance values reflect Benjamini–Hochberg–adjusted p-values. (**j**) Summary table of total transcripts belonging to IgG LC and HC, reported as the min, median and max percentage of transcripts. (**k**) SEC-seq quantified secreted IgG counts per cell, (**l**) IgG LC transcript levels per cell, and (**m**) IgG HC transcript levels per cell across the 7 cell line samples.

**Supplementary Figure 2.**
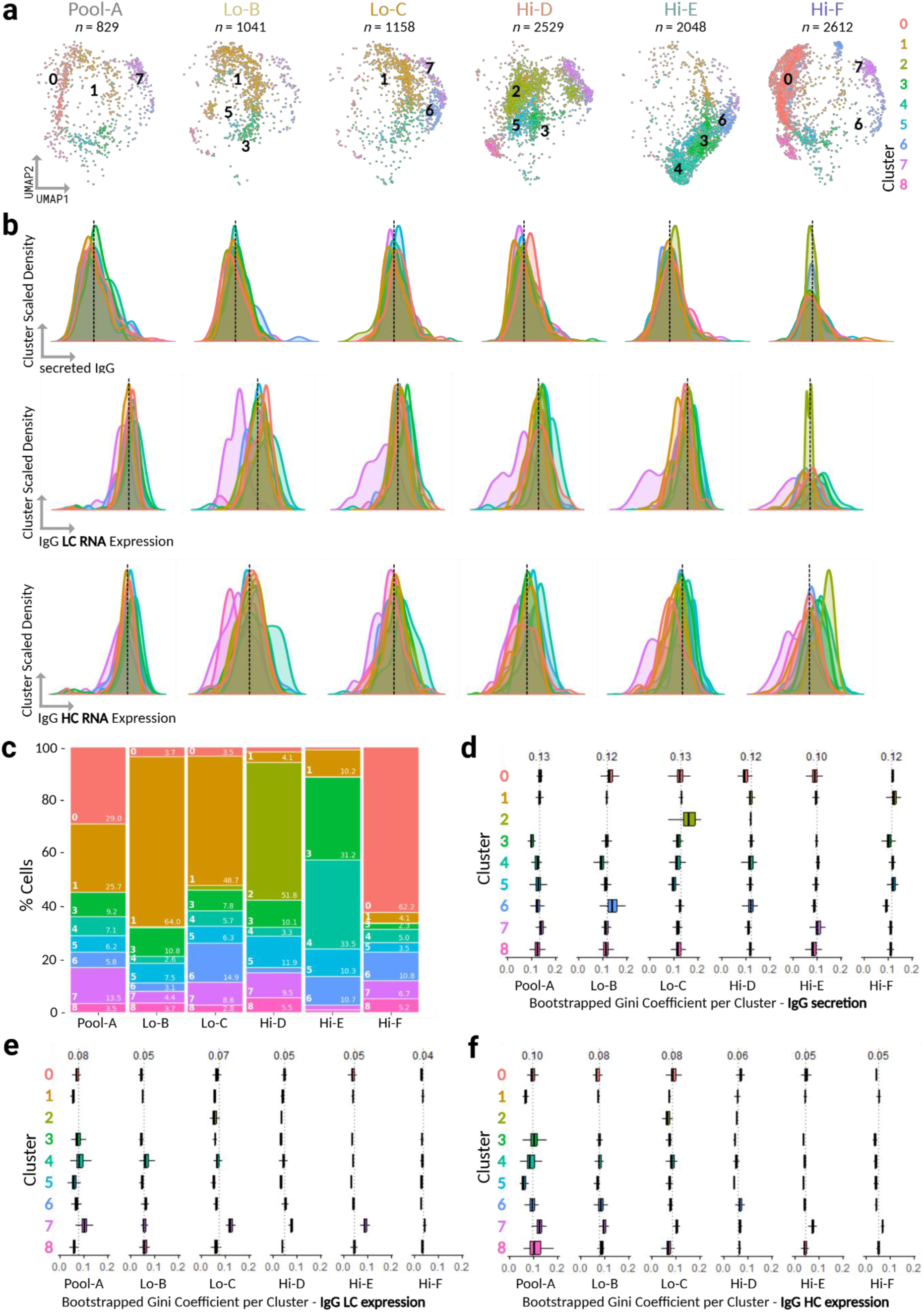
Heterogeneity of IgG RNA and protein across transcriptomic clusters and CHO cell lines using SEC-seq. (**a**) UMAP split by cell line sample in each panel colored by Louvain clusters with the top 3 abundant clusters labelled. “Lo-” indicates a low q_p_ clone and “Hi-“ indicates a high q_p_ clone. Number indicates the number of single cells (loaded in nanovial) per cell line sample. (**b**) Density plots of IgG secretion, LC expression, and HC expression per cluster per cell line sample, with median cell line sample IgG (dotted line) and median bootstrapped Gini coefficient (*G*) annotated. **(c)** Proportion of each transcriptomic cluster represented per sample. Transcriptomic cluster and proportion (%) are annotated at the bottom left and right of each color bar, respectively. **(d–f)** Bootstrapped Gini coefficients for secreted IgG (d), IgG LC (e), and IgG HC (f), shown per sample-cluster. Median cluster-level bootstrapped Gini coefficients are annotated.

**Supplementary Figure 3.**
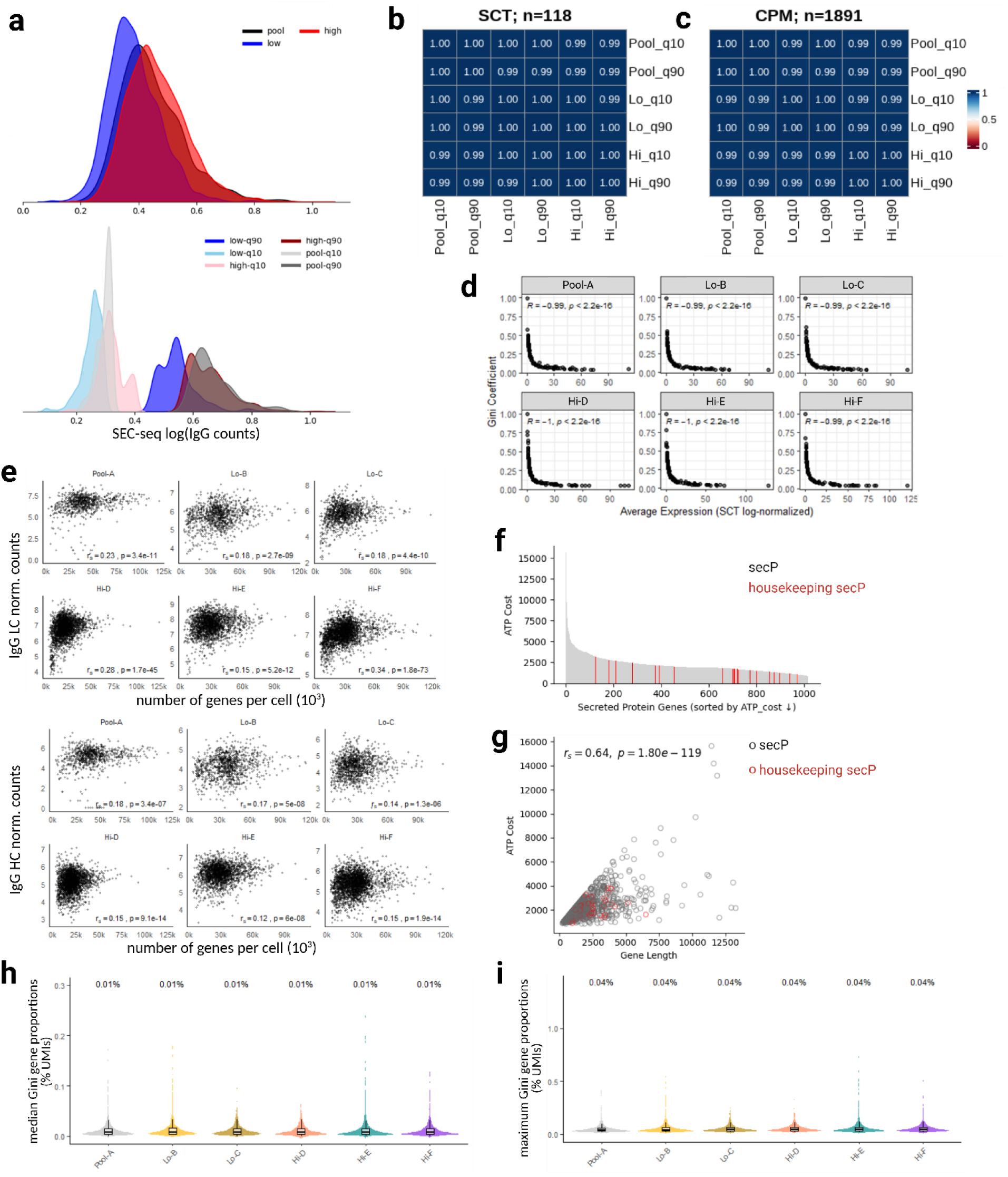
CHO housekeeping and secreted protein characteristics. (**a**) Secreted IgG profiles for parental pool (Pool-A), low secreting clones (low: Lo-B/C) and high secreting clones (high: Hi-D/E/F). Secreted IgG profiles for low (q10 = 10th percentile of IgG secretion) and high (q90 = 90th percentile of IgG secretion) IgG secreting subpopulations within each cell line group. (**b,c**) Pearson correlation matrices for average CHO housekeeping gene expression in low (q10) and high (q90) IgG secreting subpopulations using (**b**) SCTransform normalized data and (c) counts per million (CPM) normalized data. The coverage of CHO housekeeping genes per correlation matrix is indicated. Pearson correlation coefficient is represented by the colorbar gradient. (**d**) Higher average housekeeping gene expression is associated with lower Gini Coefficients, consistent with previous studies^23^. (**e**) IgG light chain (LC) and heavy chain (HC) expression demonstrate significant Spearman correlations (*r_s_*) with sequencing depth, supporting use of single-cell depth-controlled partial correlations for housekeeping gene analysis with IgG RNA levels. (**f**) ATP cost for CHO secreted proteins (secP) sorted by decreasing cost^24^, with housekeeping genes highlighted in red^23^. (**g**) ATP cost increases with gene length. Scatterplot of secP gene length versus ATP cost, with housekeeping secPs annotated in red. All significance values reflect Benjamini–Hochberg–adjusted p-values. (**h,i**) Distribution of per-gene transcriptome fractions for the top 20% most highly expressed housekeeping genes across samples (≥80th percentile by mean CPM across single cells), illustrating the limited transcriptome allocation of individual housekeeping genes relative to recombinant IgG expression. Violin plots show per-gene (**h**) medians and (**i**) maxima computed across single cells.

**Supplementary Figure 4.**
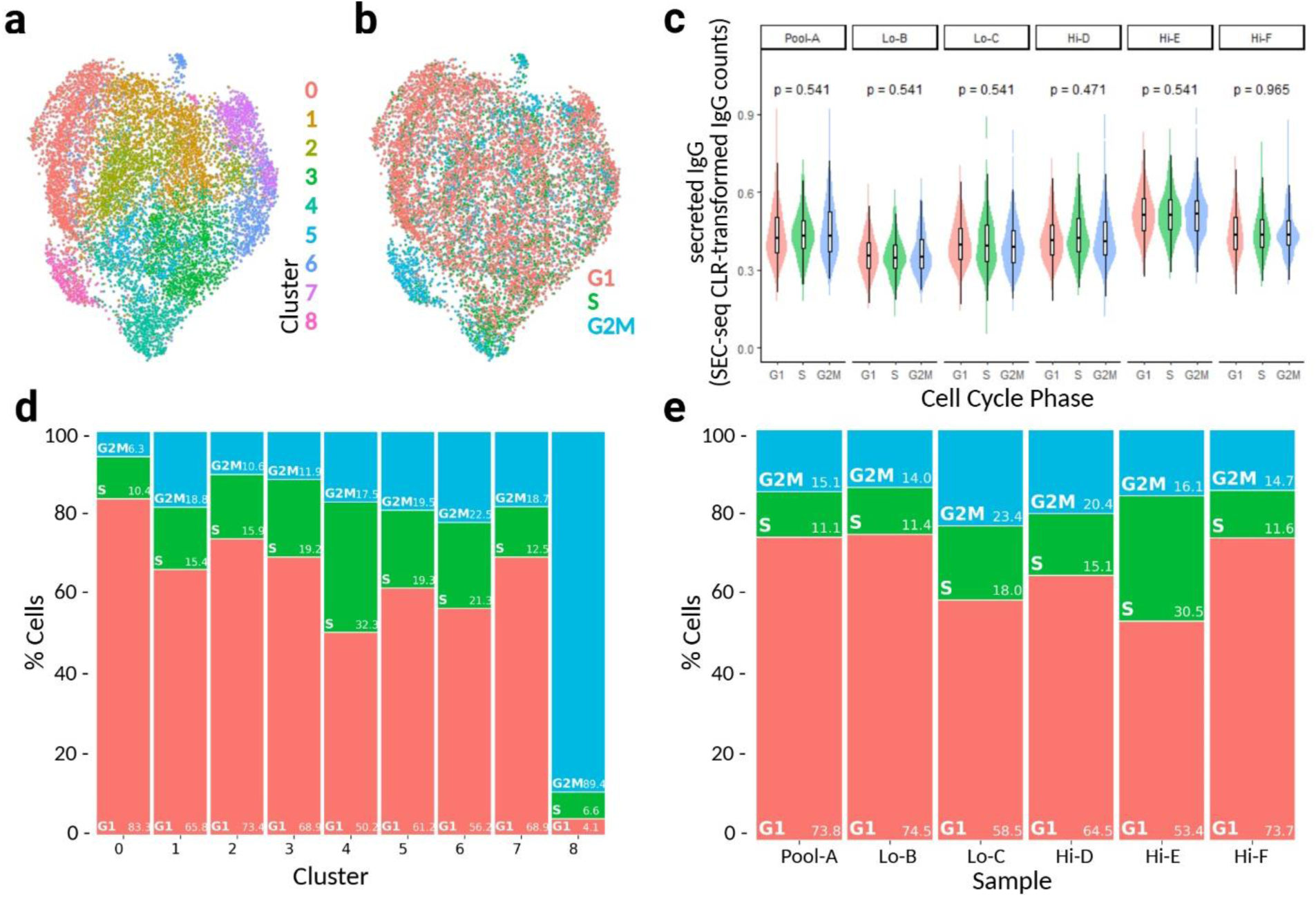
Cell-cycle composition across cell line samples and transcriptomic. (**a,b**) UMAP overlaid with (**a**) transcriptomic Louvain clusters and (**b**) cell-cycle phase after regression during SCTransform normalization. (**c**) Distribution of secreted IgG level per cell cycle phase for each cell line sample. ANOVA with BH correction indicates no significant differences in secreted IgG levels across cell-cycle phases (*p*>0.05). (**d**) Cell-cycle composition for each transcriptomic cluster. Cell-cycle phase and proportion of single-cells (%) are annotated at the bottom left and right of each color bar, respectively. (**e**) Cell-cycle phase composition for each cell line sample. Cell cycle phase and proportion of single cells (%) are annotated at the bottom left and right of each color bar, respectively.

**Supplementary Figure 5.**
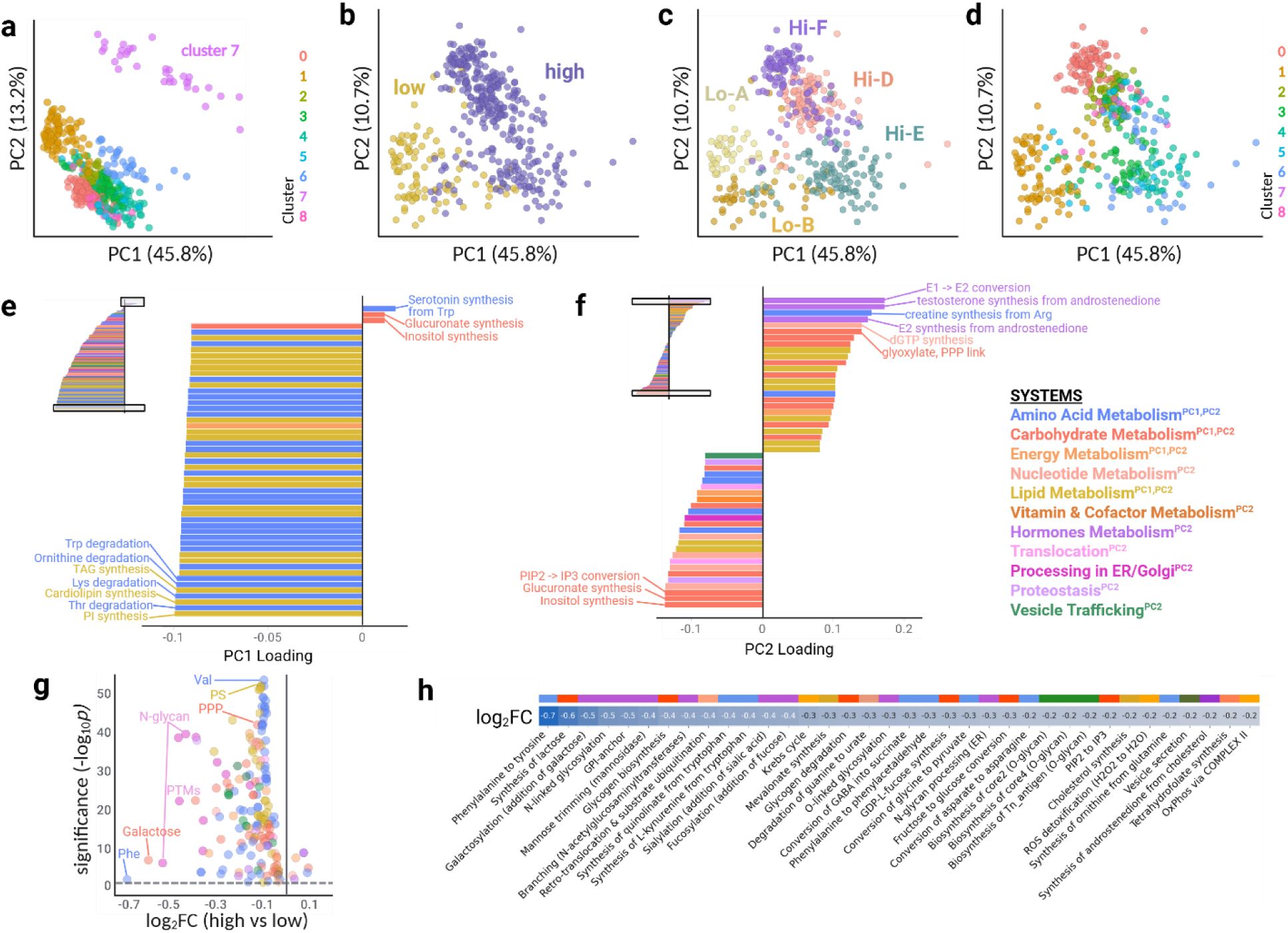
Metabolic and secretory task activity inference using scCellFie. (**a**) PCA of scCellFie metabolic-secretory task scores for pseudobulked average cell groups for each clone-cluster, overlaid with cluster. PC1 and PC2 represent the major axes of variation, with the respective percentage variation explained labeled. As cluster 7 exhibited an outlier metabolic phenotype, it was omitted from downstream analysis (**b-h**) to further find clone-cluster and high vs low differences. (**b-d**) PCA of scCellFie metabolic-secretory task scores for pseudobulked average cell groups for each clone-cluster overlaid with (**b**) low- and high-secreting phenotype, (**c**) clone, and (**d**) cluster. (e) PC1 loadings for each metabolic-secretory task colored by system annotation. The top 50 loadings (boxed in original loading plot) were highlighted, with the top 10 loaded tasks annotated. (f) PC2 loadings were similarly represented. Systems legend on the right outlines which systems contain the top 10 loadings per PC. (**g**) Volcano plot for significant differential task activities (adjusted *p*<0.05) for high- vs low-secreting clone pseudobulk groups overlaid with metabolic-secretory system. The top 4 differentially represented subsystems (for decreasing log_2_FC and decreasing significance) are annotated. (**h**) log_2_FC heatmap of differentially active metabolic tasks (|log_2_FC| > 0.2, adjusted *p*<0.05) with systems from (**f**) annotated.

**Supplementary Figure 6.**
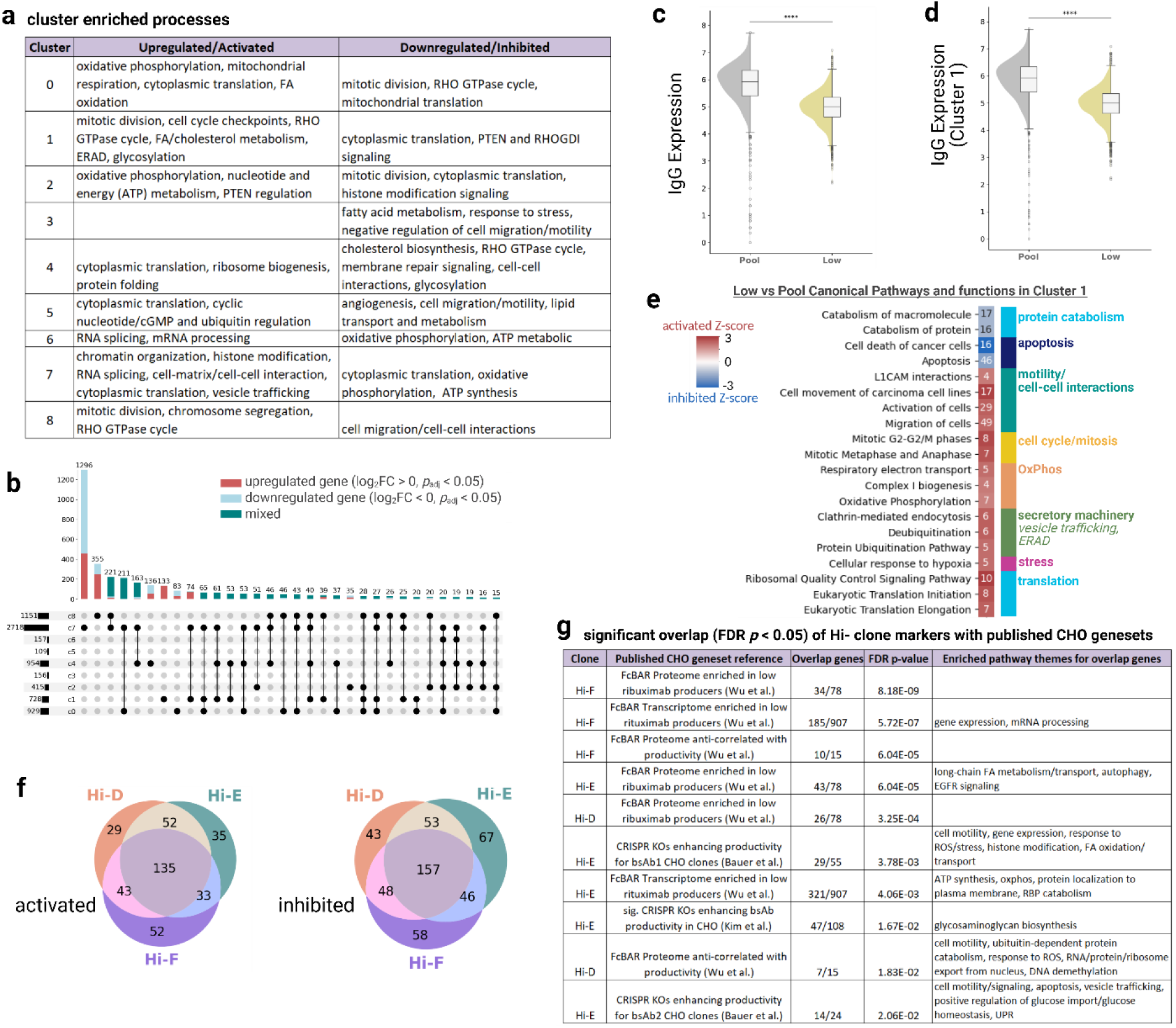
Transcriptomic cluster signatures, pool versus low-secreting clone comparisons, and overlap of downregulated high producing clonal markers with published CHO studies. (**a**) Summary of cluster-enriched biological processes derived from IPA Canonical Pathways (activated: Z-score ≥ 1.5, adjusted *p* < 0.05; inhibited: Z-score ≤ −1.5, adjusted *p* < 0.05) and GO-BP terms associated with upregulated (log_2_FC > 0, adjusted *p* < 0.05) and downregulated (log_2_FC < 0, adjusted *p* < 0.05) markers identified by Seurat *FindAllMarkers*. Cluster 3 upregulated markers did not result in any enriched biological processes. **(b)** UpSet plot showing overlap of cluster marker sets (upregulated, downregulated, or mixed). Only sets containing ≥15 genes are shown **(d)** Single-cell IgG mRNA expression (mean LC + HC) for the parental pool (Pool-A) and low-secreting clones (Lo-B/C). Mann–Whitney U one-sided tests assess greater IgG expression in the pool relative to low-secreting clones (\*\*\*\**p*<0.00001). **(e)** Same as (d), restricted to transcriptomic cluster 1, evaluated using Mann–Whitney U one-sided tests. **(f)** Heatmap of IPA-enriched Canonical Pathways and cellular functions activated (Z-score ≥ 1.5, adjusted *p* < 0.05) or inhibited (Z-score ≤ −1.5, adjusted *p* < 0.05) in low-secreting clones (Lo-B/C) relative to the parental pool within cluster 1. Color gradients denote pathway Z-scores, with annotations indicating the number of differentially expressed genes mapped to each pathway. Major pathway themes are shown in the row color bar. All significance values reflect Benjamini–Hochberg–adjusted p-values. **(f)** Comparison of activated (Z-score ≥ 1.5, adjusted p < 0.05) and inhibited (Z-score ≤ −1.5, adjusted *p* < 0.05) upstream regulators enriched in the three high-secreting clones based on IPA analysis. (**g**) Significant overlaps (hypergeometric test, adjusted *p* < 0.05) for high secreting clone markers anti-correlated with IgG relative Lo clones with published CHO enriched genesets. Overlap genes specify number of genes shared between Hi- clone markers and published geneset out of the total represented published geneset in our transcriptome background. Lack of significant pathway enrichments were left blank (adjusted *p* > 0.05). Published genesets were results from FcBAR proteome, transcriptome, and regression analysis (all adjusted *p* < 0.05) of high and low rituximab producing CHO clones (Wu et al.^28^), enriched CRISPR KOs enhancing productivity (>5% increase in normalized titer) for 2 bispecific antibody (bsAb)-producing CHO cell lines (Bauer et al.^9^), and significant CRISPR Kos enriched for bsAb producing CHO cells (Kim et al.^29^)

